# A hetero-oligomeric remorin-receptor complex regulates plant development

**DOI:** 10.1101/2021.01.28.428596

**Authors:** Nikolaj B. Abel, Corinna A. Buschle, Casandra Hernandez-Ryes, Sandy S. Burkart, Anne-Flore Deroubaix, Julia Mergner, Julien Gronnier, Iris K. Jarsch, Jessica Folgmann, Karl Heinz Braun, Emmanuelle Bayer, Véronique Germain, Paul Derbyshire, Frank L.H. Menke, Birgit Kemmerling, Cyril Zipfel, Bernhard Küster, Sébastien Mongrand, Macarena Marín, Thomas Ott

**Affiliations:** Faculty of Biology, University of Freiburg, 79104 Freiburg, Germany; Faculty of Biology, University of Munich (LMU), 82152 Planegg-Martinsried, Germany; CIBSS – Centre for Integrative Biological Signalling Studies, University of Freiburg, 79104 Freiburg, Germany; Laboratoire de Biogenèse Membranaire, Unité Mixte de Recherches UMR 5200, CNRS – Université de Bordeaux, 33140 Villenave d’Ornon, France; Technical University of Munich, Proteomics and Bioanalytics, 85354 Freising, Germany; Department of Plant and Microbial Biology, Zürich-Basel Plant Science Center, University of Zürich, 8008 Zürich, Switzerland; The Sainsbury Laboratory, University of East Anglia, Norwich Research Park, NR4 7UH, Norwich, United Kingdom; ZMBP – Center for Plant Molecular Biology, University of Tübingen, 72076 Tübingen, Germany; Technical University of Munich, Bavarian Center for Biomolecular Mass Spectrometry, 85354 Freising, Germany; Aarhus University, 8000 Aarhus, Denmark; University Hospital, LMU Munich, 80337 Munich, Germany; German Cancer Research Center (DKFZ), 69120 Heidelberg, Germany; Bavarian Center for Biomolecular Mass Spectrometry at Klinikum rechts der Isar, 81765 Munich, Germany

**Keywords:** remorin, receptor-like kinase, membrane nanodomains, plant development

## Abstract

Plant growth and development are modulated by both biotic and abiotic stress. Increasing evidence suggests that cellular integration of the corresponding signals occurs within preformed hubs at the plasma membrane called nanodomains. These membrane sub-compartments are organized by multivalent molecular scaffold proteins, such as remorins. Here, we demonstrate that Group 1 remorins form a hetero-oligomeric complex at the plasma membrane. While these remorins are functionally redundant for some pathways their multivalency also allows the recruitment of specific interaction partners. One of them, the receptor-like kinase REMORIN-INTERACTING RECEPTOR 1 (RIR1), that acts redundantly with the closely related receptor NILR2, is specifically recruited by REM1.2 in a phosphorylation-dependent manner. Overlapping developmental phenotypes suggest that the REM/RIR complex regulates key developmental pathways.

## Introduction

Being exposed to a highly dynamic environment, organisms have to simultaneously integrate a plethora of extracellular stimuli. In plants, the cell wall and the plasma membrane (PM) form crucial barriers to compartmentalize, protect and modulate cytosolic processes. PM functionality depends on multiprotein entities that can form higher order assemblies in the nanometer (nanodomains) or micrometer (microdomains) range (Ott, 2017; Jaillais and Ott, 2019). Among others, membrane-resident receptors have been reported to localizes to such nanodomains ((Haney et al., 2011; Bücherl et al., 2017; Burkart and Stahl, 2017; Liang et al., 2018; Gronnier et al., 2020)). Using fluorescently-tagged fusion proteins, membrane nanodomains appear as punctate structures at the cell surface with a low degree of lateral mobility (Ott, 2017). Although precise molecular functions and mechanisms of receptor nanoclustering have only been unraveled for few cases so far (Gui et al., 2016; Liang et al., 2018) it is widely believed that these structures provide organizational units that allow protein complex formation and stabilization in complex cellular environments. Given the additional complexity of extracellular stimuli that need to be perceived and translated to generate specific cellular outputs and the comparably limited availability of multivalent proteins that could serve such functions, it can be postulated that hetero-oligomeric scaffolds such as remorin protein may act as hubs for signal integration (Jaillais and Ott, 2019).

Remorins form a multigene family with 16 members belonging to 5 different subgroups in *Arabidopsis thaliana* while legumes evolved an additional group (group 2) (Raffaele et al., 2007). In general, remorins are comprised of a highly diverse, N-terminal intrinsically disordered region (IDR) and a conserved coiled-coil domain-containing C-terminal segment (Raffaele et al., 2007). The latter harbors a PM binding motif (RemCA) that anchors remorins to the cytosolic leaflet of the PM, a process that is further supported by palmitoylation (Raffaele et al., 2009; Perraki et al., 2012; Konrad et al., 2014; Gronnier et al., 2017; Legrand et al., 2019). Although the molecular function remains to be elucidated for most of these proteins, recent work showed that SYMBIOTIC REMORIN 1 (SYMREM1), a group 2 remorin from *Medicago truncatula*, interacts with symbiotic RLKs and controls rhizobial infections by recruiting and immobilizing the LYSIN MOTIF RECEPTOR-LIKE KINASE 3 (LYK3) in specific membrane nanodomains (Lefebvre et al., 2010; Liang et al., 2018). Additional evidence for remorin-mediated modulation of receptor complexes has also been presented in rice, where the group 4 remorin OsREM4.1 regulates a phytohormonal cross-talk by reversibly associating with an RLK complex in a phosphorylation-dependent manner (Gui et al., 2016). The most abundant remorins, however, belong to Group 1, with two members (REM1.2 and REM1.3) being among the most highly expressed proteins (Top 20%) in Arabidopsis (Mergner et al., 2020). Even though an involvement of Group 1 remorins in phytohormone controlled processes has been suggested (Alliotte et al., 1989; Yamada et al., 1998; Demir et al., 2013; Gui et al., 2016; Huang et al., 2019), the molecular function of these proteins remains elusive. Several studies have independently demonstrated that Group 1 remorins regulate viral spreading (Raffaele et al., 2009; Son et al., 2015; Perraki et al., 2018; Cheng et al., 2020), possibly by either regulating plasmodesmatal (PD) conductance (Perraki et al., 2014; Gronnier et al., 2017; Huang et al., 2019) and/or even PD biogenesis (Wei et al., 2020). However, considering that the subcellular localization of Group 1 remorins is not limited to PDs (Raffaele et al., 2009; Jarsch et al., 2014), it appears likely that they play additional roles. Their ability to form higher order homooligomeric structures (Bariola et al., 2004; Marin et al., 2012; Legrand et al., 2019; Martinez et al., 2019) and the presence of an IDR (Marin et al., 2012) make them prominent candidates to function as multivalent scaffolds.

As such signalling hubs would be central for plant survival or at least fitness of the individuum in fluctuating environments, they can be protected by functional redundancy. To address this, we genetically analysed the hetero-oligomeric Group 1 remorin complex at the plasma membrane and demonstrate that specificity can be generated within the complex by individual members despite their overall functional redundancy. Functional specification is exemplified by the identification of the redundant receptor pair RIR1/NILR2 out of which RIR1 specifically associates with REM1.2, while NILR2 did not interact with group 1 remorins.

## Results

### Genetic redundancy within the remorin Group 1 family

Two members of the Group 1 sub-clade, REM1.2 (At3g61260) and REM1.3 (At2g45820), belong to the top 20% most abundant proteins in Arabidopsis (Mergner et al., 2020). While average protein levels in rosette leaves for REM1.4 (At5g23750) are almost two orders of magnitude lower compared to REM1.2 and REM1.3, the fourth Group 1 protein (REM1.1; At3g48940) was only weakly detected (in few samples and close to the detection limit) by mass spectrometry. However, low abundant transcripts of *REM1.1* have been reported (Jarsch et al., 2014).

To genetically dissect functional redundancy and specificity within Group 1 remorins we used the already described *rem1.2-1* and *rem1.3-2* single mutant lines (Jarsch et al., 2014) and created a *rem1.2/rem1.3* double and a *rem1.2/1.3/1.4* triple mutant by introgressing the *rem1.4-3* (N651091) allele. No knock-out allele was identified for REM1.1. None of these lines showed a significant developmental phenotype when scoring rosette diameter over a period of 28 days (Fig. 1A). As this was in contrast to previous reports (Huang et al., 2019; Wei et al., 2020), we tested the published *rem1.2/1.3c-L27* knock-out line, but did also not observe the reported growth defect under our growth conditions (Fig. S1A), while the inducible over-expressor XVE:REM1.2 was indeed growth-retarded as previously reported (Huang et al., 2019) (Fig. S1B). This indicates that the *rem1.2/1.3* developmental phenotype might be conditional. Thus, we tested viral spreading as this has been reported to be altered in Group 1 remorin mutants (Raffaele et al., 2009; Perraki et al., 2018). For this, *rem1.2/1.3* and *rem1.2/1.3/1.4* mutants were infected with a fluorescently labelled *Plantago asiatica* mosaic virus (PlAMV), a Potexvirus that has previously been shown to infect *A. thaliana* (Minato et al., 2014; Hashimoto et al., 2016). Virus propagation was scored 7 dpi by determining the size of the PlAMV-GFP foci. Wild-type Col-0 plants showed average infection sites of 0.58 mm^2^ (Fig. 1B). While areas of PlAMV infection foci in *rem1.2* and *rem1.3* single mutants were indistinguishable from those found on WT plants, the *rem1.2/1.3* double mutant was significantly more susceptible to PlAMV. This effect was further pronounced in the triple mutant (Fig. 1B). These data demonstrate that REM1.2, REM1.3 and REM1.4 act redundantly during viral spreading.

**Fig. 1:**
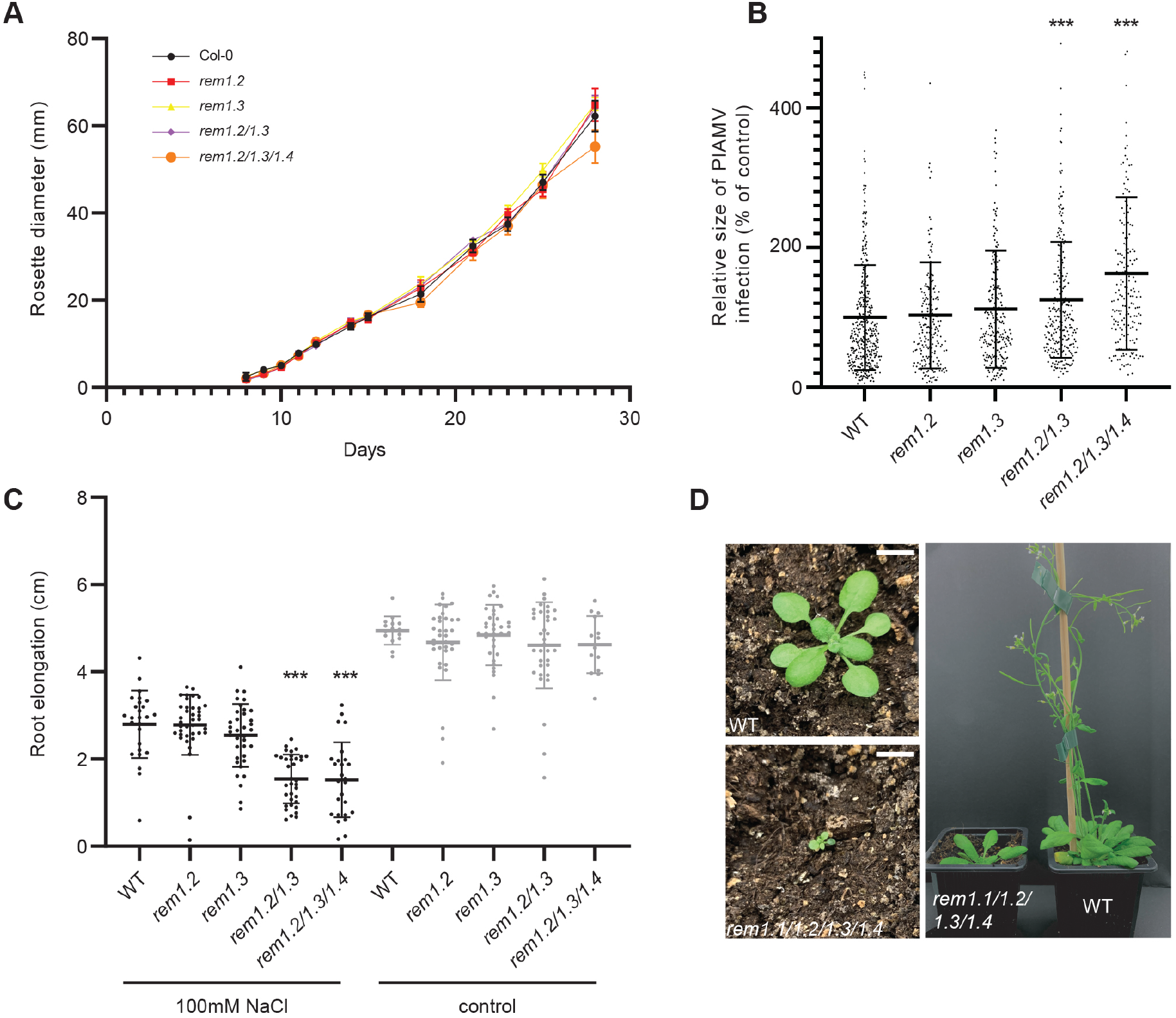
Group 1 remorins are functionally redundant and essential for plant growth. (A) No growth phenotype was observed in *rem1.2* (red), *rem1.3* (yellow) and *rem1.2/1.3* (purple) mutants when scoring rosette diameter over time. (B) Assessing spreading of the *Plantago asiatica* mosaic virus (PlAMV) revealed an increased susceptibility of *rem1.2/1.3* double and *rem1.2/1.3/1.4* triple mutants. (C) Plants were grown on 100mM NaCl for seven days before root length was scored. While no differences were found in the controls (mock), growth of *rem1.2/1.3* double and *rem1.2/1.3/1.4* triple mutants was significantly reduced under salt stress. (D). A *rem1.1^HET^/1.2^HOM^/1.3^HOM^ /1.4^HOM^* quadruple mutant generated by CRISPR-CAS9 showed a dwarf phenotype and the inability to set seeds. Scale bars represent 1cm. Significance was tested using a Dunnett’s multiple comparisons test with p<0.001 (***).

To test whether this phenomenon can also be observed for other cellular processes we made use of the fact that heterologous over-expression of the Group 1 remorin SiREM6 from foxtail millet (*Setaria italica*) in *A. thaliana* resulted in increased salt tolerance of the transgenic lines (Yue et al., 2014). Therefore, we germinated seeds on 0.5x MS plates, before we transferred seedlings onto 100 mM NaCl for seven days and scored their root lengths. While no significant differences were observed between WT and the *rem1.2* and *rem1.3* single mutants, the *rem1.2/1.3* double and *rem1.2/1.3/1.4* triple mutant showed a highly significant reduction in root length under salt stress (Fig. 1C). Thus, we conclude that all three REMs act redundantly in the same pathway with respect to salt tolerance.

To fully assess functional redundancy within this remorin clade, we generated quadruple *rem1.1/1.2/1.3/1.4* mutant lines by CRISPR-Cas9. For this, we designed guide RNAs targeting REM1.1 with REM1.2 and REM1.4 as putative off-targets. While we were never able to generate homozygous T2 lines, we were able to rescue two lines in the T1 generation being heterozygous in the *REM1.1* locus of which all individuals showed a dwarf phenotype when grown on soil (Fig. 1D). This unequivocally demonstrates that Group 1 remorins, which act at least partially redundant, are essential for plant growth but may exhibit their full functional potential under more natural conditions that require multi-layered integration of environmental cues.

### Remorins form a heteromeric and multivalent scaffolding complex at the plasma membrane

Given their functional redundancy and the fact that remorins have already been described to form homo-oligomeric complexes (Bariola et al., 2004; Marin et al., 2012; Perraki et al., 2012; McBride et al., 2017; Martinez et al., 2019), we asked whether the highly abundant REM1.2 and REM1.3 proteins also form hetero-oligomeric complexes in *A. thaliana*. For this, we immunoprecipitated both fluorophore-tagged remorins expressed in their corresponding mutant backgrounds and under control of their native promoters using GFP-nanotraps and confirmed association of the endogenous REM1.2 and REM1.3 using a specific antibody against the endogenous proteins (Fig. 2A). To obtain a more global view and to dissect whether we can detect specific interaction profiles for both remorins, we immunoprecipitated REM1.2 and REM1.3 and determined associated proteins by untargeted mass-spectrometry. We obtained high confident spectra representing 2163 and 2140 unique protein identifiers found in association with REM1.2 and REM1.3, respectively with 2108 protein identifiers being identical for both remorins. However, despite this large overlap, both genotypes could clearly be separated based on their protein intensity profiles (Fig. 2B) and significant different protein interaction partners (Fig. 2C). These data support the observed genetic redundancy but also imply that specificity within the hetero-oligomeric REM complex may be generated by individual members.

**Fig. 2:**
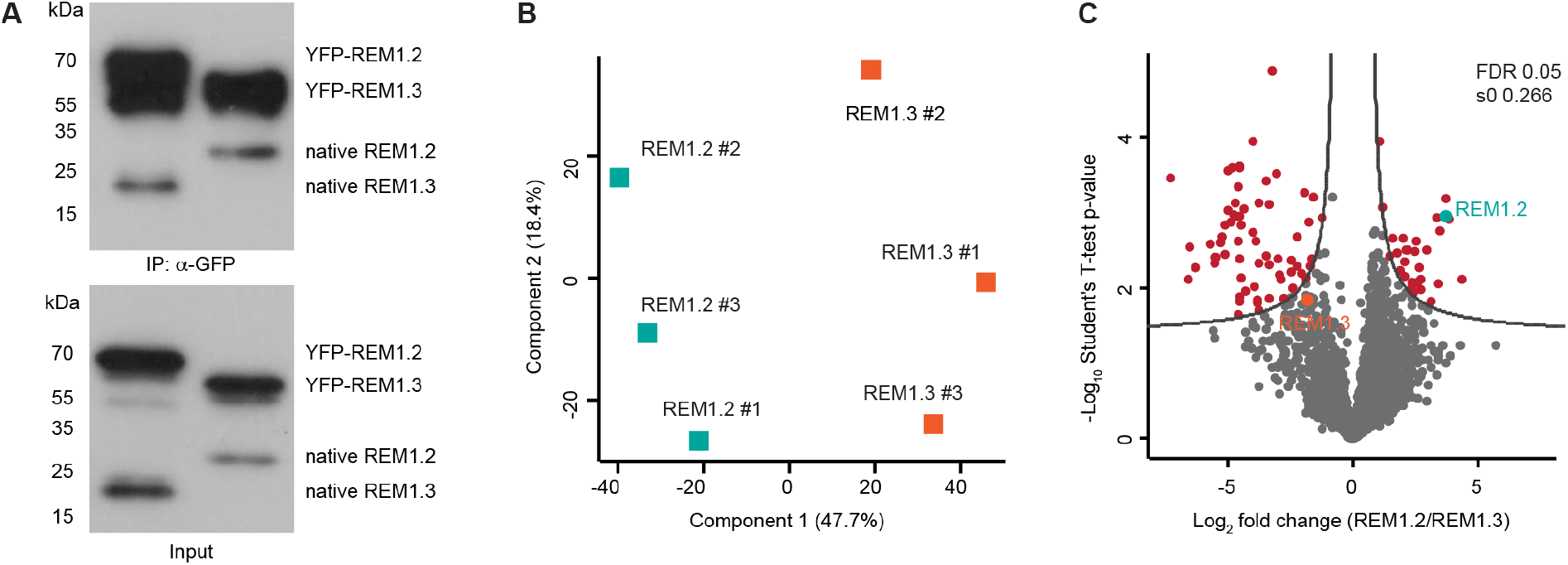
REM1.2 and REM1.3 form a hetero-oligomeric complex in planta. (A) Co-immunoprecipitation of REM1.3 and REM1.2 in complemented *rem1.2* and *rem1.3* mutant lines, respectively, using GFP nanotraps and detecting endogenous REM1.2 and REM1.3 as well as the YFP fusion proteins by a specific antibody recognizing both proteins. (B) Principal component analysis (PCA) of REM1.2 and REM1.3 immuno-precipitation proteome analysis. (C) Volcano plot of REM1.2 and REM1.3 immuno-precipitation proteome fold changes and p-values. Significant different abundant proteins are depicted in red (Student’s t-test, FDR < 0.05). REM1.2 and REM1.3 are depicted in blue and orange, respectively.

### The membrane-resident LRR-malectin receptor RIR1 interacts with REM1.2

Since the representation of receptors was very low in our proteome data set, aiming to identify new receptors that regulate plant growth and assuming that several of these RLKs are difficult to extract and often lowly abundant, we decided to supplement our interaction data with a targeted screen where we tested for pairwise interactions between REM1.2 and the cytoplasmic domains (CDs) of 55 different RLKs by conducting a classical yeast-2-hybrid screen (Table S1). These RLKs had originally be selected to their putative involvement in plant-microbe interactions. Using REM1.2 as bait, a single CD encoded by At1g53440 was found to reproducibly interact with this remorin protein. Growth of yeast colonies on triple selective – LWH drop out medium was sustained in all three tested dilutions and maintained in the presence of 2.5 mM 3-Amino-1,2,4-triazole (3-AT), a competitive inhibitor of histidine biosynthesis (Fig. 3A). Therefore, we named the corresponding protein REMORIN-INTERACTING RECEPTOR 1 (RIR1). In this experimental system, the RIR1/REM1.2 interaction was found to be highly specific as the RIR1 CD was unable to associate with REM1.3 (Fig. 3A). Functionality and stability of both remorin clones were independently verified by testing for homo-oligomerisation (Fig. 3A) and detection of full-length proteins by Western blotting (Fig. S2).

**Fig. 3:**
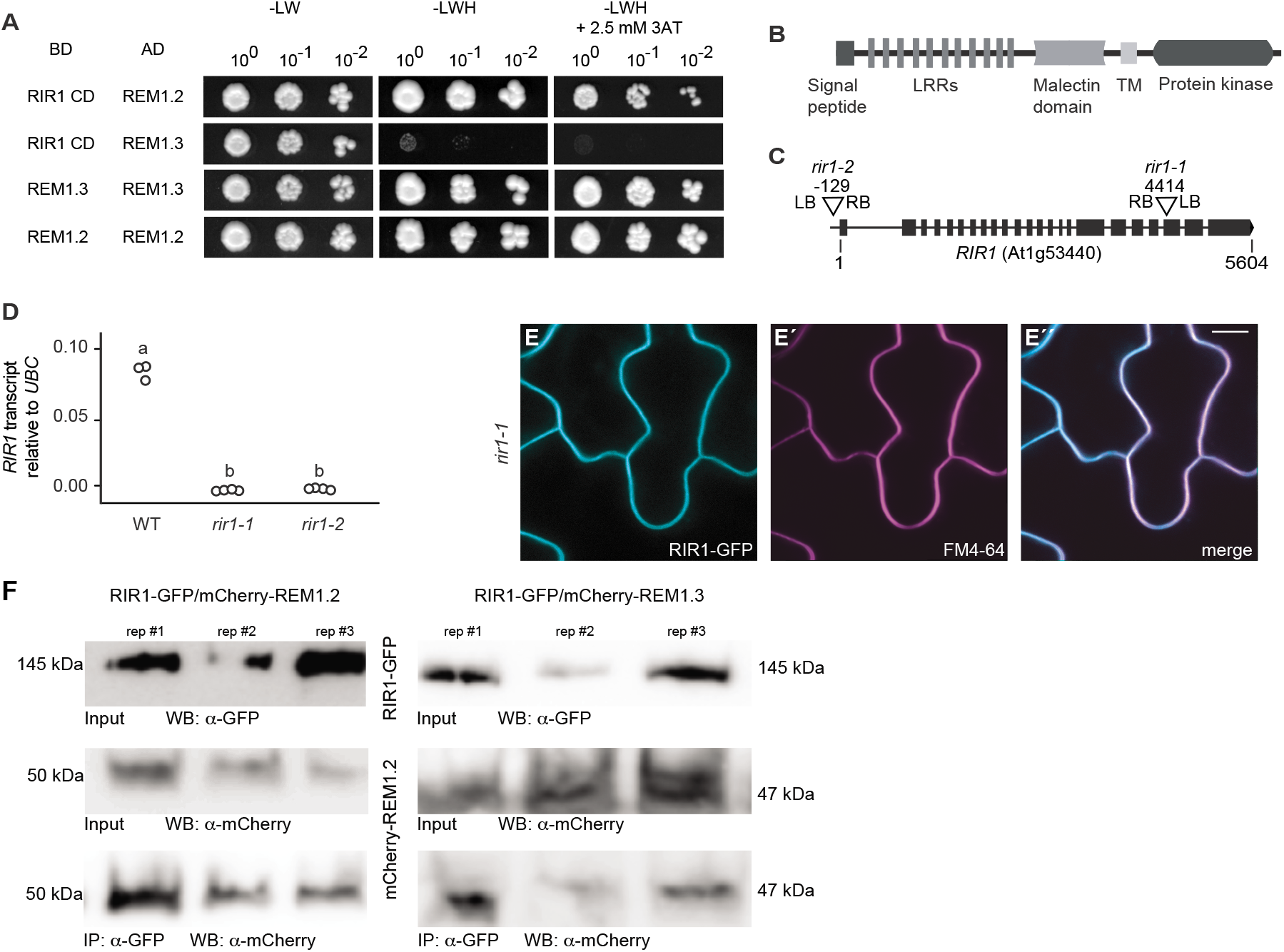
Identification of the REMORIN-INTERACTING-RECEPTOR 1 (RIR1). (A) Screening for interaction partners for REM1.2 by independently co-expressing 55 isolated cytoplasmatic domains (CDs) of selected receptor-like kinases yielded in a strong interaction between RIR1 (At1g53440) with REM1.2 but not with REM1.3. Both remorin were able to homo-dimerize in yeast, demonstrating functionality of the clones. (B) Schematic illustration of the RIR1 protein structure with 11 leucin-rich repeats (LRRs) followed by a malectin domain, a transmembrane domain (TM) and a protein kinase domain. (C) Identification of two mutant alleles in public SALK mutant collection. Numbers indicate nucleotide positions. (D) Quantitative RealTime PCR confirmed the loss of transcript in both *rir1* mutant alleles. (E) Expressing of a fluorophore-tagged RIR1 driven by the endogenous *rir1* promoter in the *rir1-1* mutant background revealed plasma membrane localization of the receptor and co-localization with the membrane dye FM4-64 in five week old leaf epidermal cells (E’-E’’). Scale bars indicate 10μm. (F) Co-immunoprecipitation of RIR1 from independent *rir1-1/ProRIR1-RIR1-GFP* lines expression either a *ProREM1.2-mCherry-REM1.2* or a *ProREM1.3-mCherry-REM1.3* construct. background done in three independent replicates. REM1.2 and REM1.3 co-immunoprecipitating with RIR1 were detected with an a-mCherry antibody.

The *RIR1* gene encodes a receptor-like kinase with 11 predicted leucine-rich repeats (LRRs) and a malectin-domain in its extracellular domain and a kinase domain in its C-terminal intracellular region (Fig. 3B). Previous phylogenetic analyses placed this protein into the subgroup VIII-2 of the LRR-RLK family (Shiu and Bleecker, 2001).

Since we were unable to identify RIR1 as a possible candidate from our initial MS data (Table S2) we aimed to verify this interaction in the Arabidopsis homologous system. For this, we isolated two independent SALK T-DNA insertion lines (*rir1-1* (Salk_130548) and *rir1-2* (Salk_057812)). Re-sequencing the insertion loci confirmed the presence of the T-DNAs at 129 base pairs (bp) upstream of the predicted translational start codon in *rir1-2* and within the 21^st^ exon at nucleotide position 4414 in *rir1-1* (Fig. 3C). Expression analysis confirmed a transcriptional knock-down for both lines (Fig. 3D). However, none of the two *rir1* mutant alleles showed a significant developmental phenotype (Fig. S3). Next, we introduced a RIR1-GFP fusion protein driven by the native *RIR1* promoter (1.8 kb upstream of the start codon; *ProRIR1:RIR1-GFP*) in the *rir1-1* mutant background. Based on fluorescent signal intensities we selected two independent complemented lines (*rir1-1*/RIR1-GFP#3 and #4) that were used in the following analyses. Subcellular localization of RIR1-GFP in these transgenics revealed exclusive fluorescence at the plasma membrane that co-localized with the styryl dye FM4-64 (Fig. 3E). To test the RIR1-remorin interaction *in planta* we transformed a *ProREM1.2-mCherry-REM1.2* and a *ProREM1.3-mCherry-REM1.3* into the *rir1-1*/RIR1-GFP#3 line. Using a GFP-nanotrap, we immunoprecipitated RIR1-GFP from these lines and detected mCherry-REM1.2 and mCherry-REM1.3 co-precipitating with RIR1 (Fig. 3F). As we could not detect any interaction between REM1.3 and RIR1 in yeast, these data indicate that REM1.3 is an integral part of the RIR1-REM1.2 complex, but interacts rather with REM1.2 than with the receptor itself. To test this *in planta*, we used cotyledons of 5 days old homozygous seedlings to perform Fluorescence Lifetime Imaging Microscopy-Foerster Resonance Energy Transfer (FLIM-FRET). FLIM analysis on the *rir1-1*/RIR1-GFP#3/mCherry-REM1.2-line (D= donor; A= acceptor) indeed revealed significantly reduced lifetimes (τ) with average values of τ_DA_ 2.21 +/-0.05 ns while the donor-only (τ_D_) lifetime of GFP in the *rir1-1*/RIR1-GFP#3 line was 2.58 +/-0.03 nanoseconds (ns) (Fig. 4). In contrast, no decrease in fluorescence lifetime was observed in *rir1-1*/RIR1-GFP/mCherry-REM1.3-line with average τDA-values of 2.62 +/-0.01 ns (Fig. 4). These data clearly support that RIR1 is recruited into the hetero-oligomeric REM1.2/REM1.3 complex where it predominantly and directly interacts with REM1.2.

**Fig. 4:**
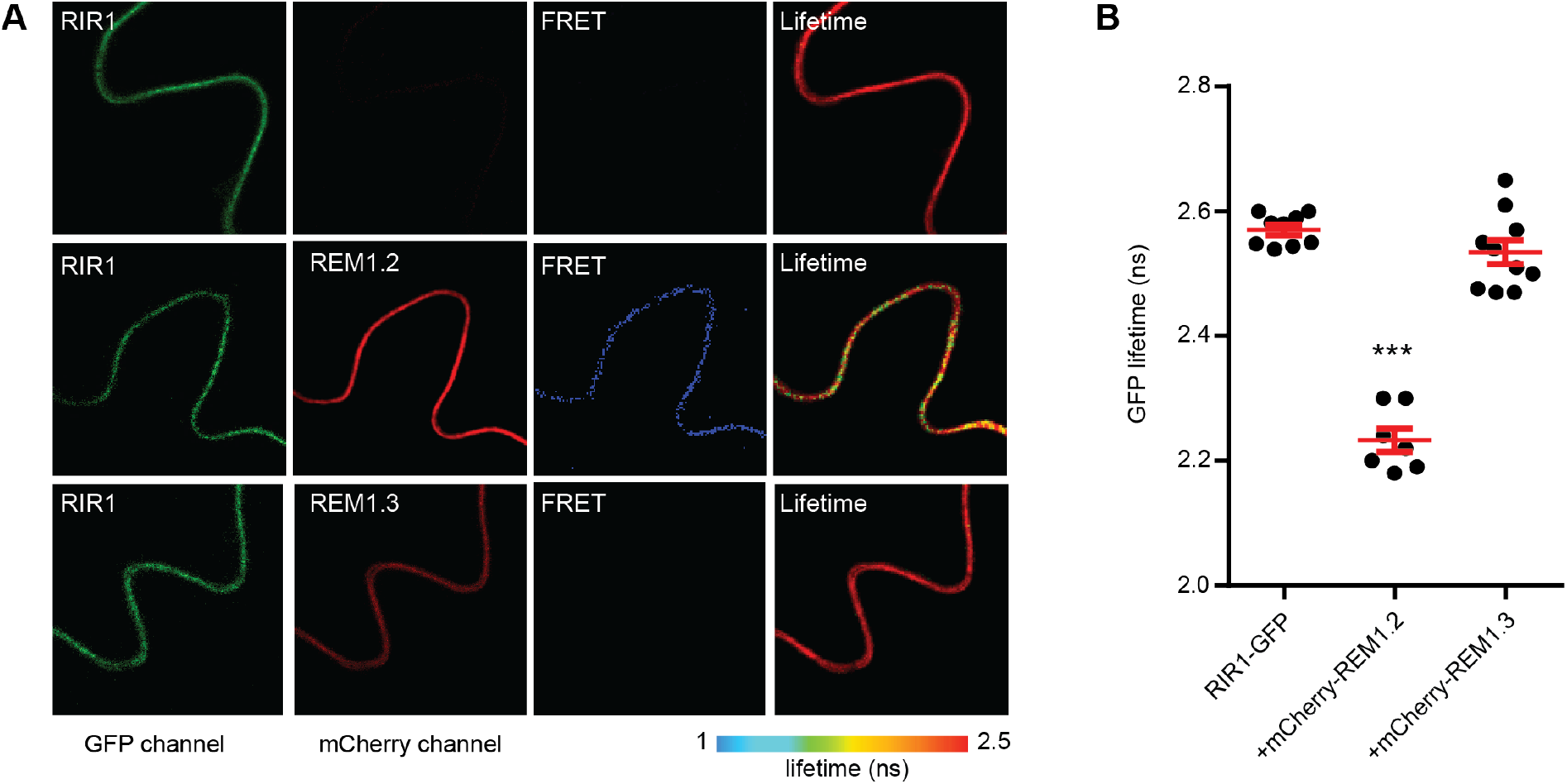
Confirming the direct interaction between REM1.2 and RIR1 in stable transgenic Arabidopsis lines. Fluorescence Lifetime Imaging Microscopy-Förster Resonance Energy Transfer (FLIM-FRET) was applied to *rir1*/RIR1-GFP, *rir1*/RIR1-GFP/mCherry-REM1.2 and *rir1*/RIR1-GFP/mCherry-REM1.3 plants. Experiments were performed on cotyledons of 5 days old seedlings. (A) Representative pictures on cells used for FRET analysis. (B) Quantification of the GFP lifetime. Significance was tested using a Dunn’s multiple comparisons test with p<0.001 (***).

### RIR1 localizes to membrane nanodomains independent of REM1.2/REM1.3

Since several RLKs have been reported to localize to distinct membrane nanodomains (Haney et al., 2011; Wang et al., 2015; Bücherl et al., 2017; Hutten et al., 2017) we asked whether RIR1 also localizes to these structures at the PM. In order to minimize background fluorescence, we applied Variable Angle Total Internal Reflection Microscopy (VA-TIRFM). Analysis of the *RIR1-1*/RIR1-GFP#3 line confirmed that RIR1 localizes to small nanodomains that appeared as punctate structures with high signal intensities at the cell surface (Fig. S4A). Single particle tracking (SPT) revealed an immobility of the RIR1-labelled nanodomains (Fig. S4B-D), which is a hallmark of these structures in plants. To assess whether the REM1.2/REM1.3 complex is required for the lateral immobilization of RIR1, we transformed the *ProRIR1:RIR1-GFP* construct into the *rem1.2* single and the *rem1.2/1.3* double mutant backgrounds. Analysis of homozygous lines showed neither an altered nanodomain localization pattern (Fig. S4A) nor any differences in RIR1 mobility (Fig. S4B-D) in these mutant backgrounds indicating that the REM1.2/REM1.3 scaffolding complex is not required for nanodomain recruitment of RIR1.

### REM1.2 and RIR1 interact in a phosphorylation-dependent manner

Since phosphorylation of Group 1 remorins has been repeatedly reported (Benschop et al., 2007; Keinath et al., 2010; Kohorn et al., 2014; Gui et al., 2016; Menz et al., 2016; Nikonorova et al., 2018; Perraki et al., 2018), we tested whether the RIR1/REM1.2 interaction depends on this post-translational modification. For this, we created a kinase-dead variant of RIR1, where we mutated the aspartate (D) in the conserved DFG motif within the activation loop of the kinase domain into an asparagine (N) (RIR1^D811N^). Mutations of the conserved aspartate are known to result in kinase-dead mutants (Yoshida and Parniske, 2005). Indeed, the D811N mutation in RIR1 entirely abolished the interaction with REM1.2 in yeast (Fig. 5A).

**Fig. 5:**
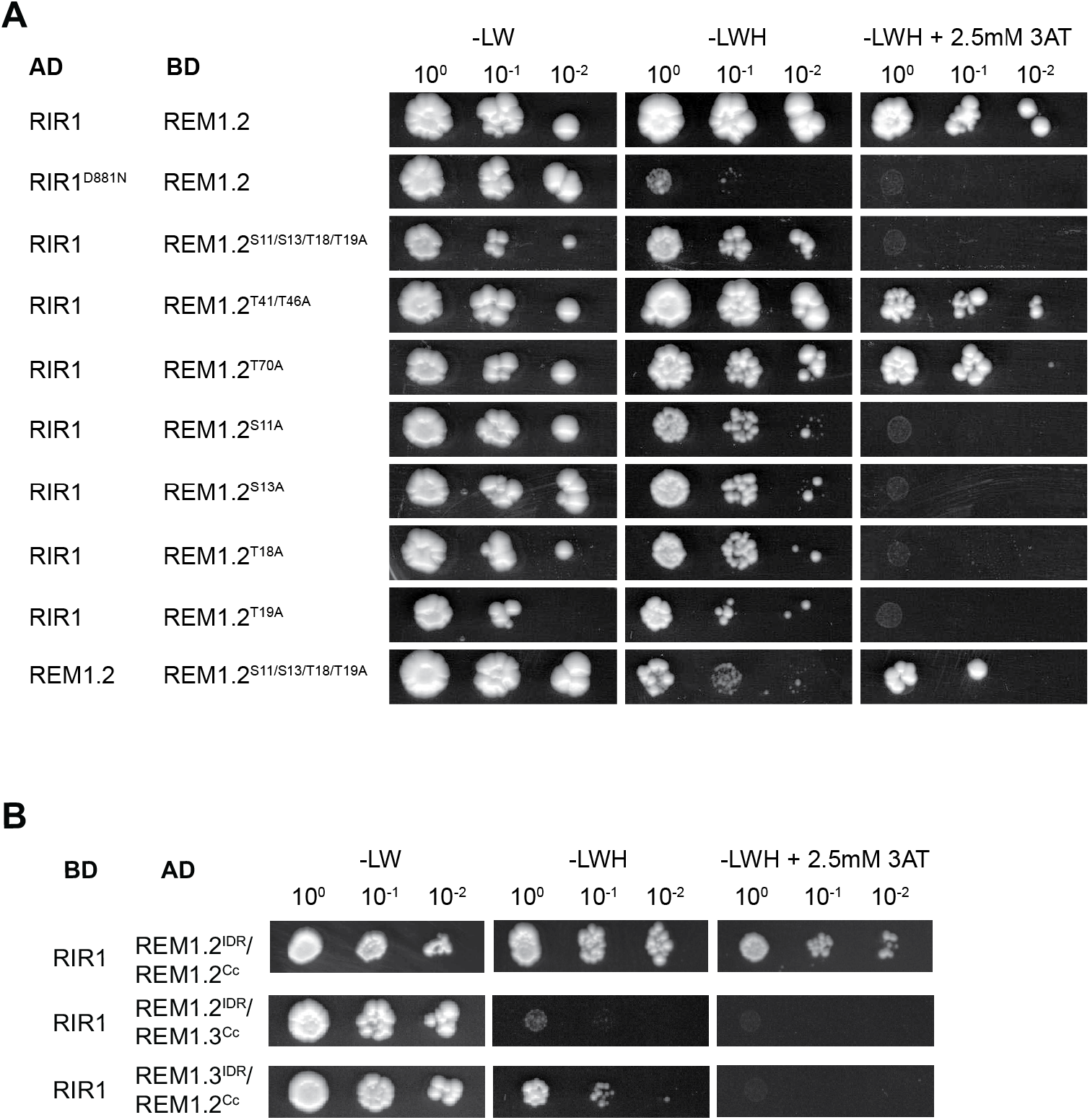
Phosphorylation-dependent interaction between RIR1 and REM1.2 in yeast. (A) GAL4 Y2H assay of RIR1 wild type or kinase-dead (D811N) with REM1.2 wild type or phospho-ablative (S11/S13/T18/T19A, S11A, S13A, T18A, T19A, T41/T46A, T70A) mutants. (B) Testing chimeric remorin proteins where the intrinsically disordered N-terminal region (IDR) and the conserved coiled coil (Cc) region were reciprocally exchanged between REM1.2 and REM1.3. Remorins were fused to the activation domain (AD) and the kinases to the binding domain (BD) of GAL4. Transformants were grown on control medium (-LW) and selective medium (-LWH +/−2.5 mM 3AT) in three consecutive dilutions. S= serine, T= threonine, L = leucine, W = tryptophan, H = histidine, 3-AT = 3-amino-1,2,4-triazole.

Since the majority of phosphorylation sites are located within the intrinsically disordered region (IDR) of Group 1 remorins (Marín and Ott, 2012), we first tested the contributions of the IDR and the structured coiled-coil C-terminal region (Cc) to RIR1/REM complex formation. Interestingly, chimeric variants of REM1.2 and REM1.3 (REM1.2^IDR^:REM1.3^Cc^ and REM1.3 ^IDR^:REM1.2^Cc^) failed to interact with RIR1 in yeast (Fig. 5B) indicating that both regions contribute to the interaction. Next, we created a series of phosphoablative mutations (REM1.2^S11/S13/T18/T19A^, REM1.2^T41/T46A^ and REM1.2^T70A^) in putative REM1.2 phospho-sites focusing on residues that are not conserved between REM1.2 and REM1.3. While the REM1.2^S11/S13/T18/T19A^ mutant failed to interact with RIR1, the REM1.2^T41/T46A^ and REM1.2^T70A^ mutants showed sustained yeast growth under selective conditions (Fig. 5A). Expression of all clones not yielding an interaction was confirmed by Western blot analysis (Fig. S5). These results indicate that the SSTT cluster between residues 10 and 20 within the REM1.2 N-terminal region contributes to the REM1.2/RIR1 complex formation.

### RIR1 is a component of a functionally redundant receptor cluster

As mentioned above, our *rir1* mutant alleles did not show any significant growth defects (Fig. S3) as demonstrated for the *rem1.1/1.2/1.3/1.4* quadruple mutant (Fig. 1D). To test for overlapping phenotypes between remorins and the RIR1 receptor, we inoculated *rir1-1* and *RIR1-2* mutants with PlAMV and scored viral infection foci after 7 days. Even though both alleles showed a slightly increased susceptibility to PlAMV, these effects were significantly smaller compared to those observed for the *rem1.2/1.3/1.4* triple mutant (Fig. 1B).

To obtain more insights into proteins interacting with RIR1, we performed immunoprecipitation followed by mass-spectrometry using the *rir1-1*/RIR1-GFP#3 line and a line expressing the membrane marker LTI6b-GFP as a control. Interestingly, we identified an additional malectin-domain containing RLKs (*At1g53430*) directly flanking the *RIR1* gene (*At1g53440*), among others, as putative interactors of RIR1 (Table S3) with At1g53430 sharing about 90% sequence similarity with RIR1. However, we identified two exclusive unique peptides in two independent biological replicates and these peptides were observed with multiple spectra with good Mascot scores. In contrast, a third receptor of this gene cluster (*At1G53420*) does not have any exclusive unique peptides and so most spectra are likely to derive from either RIR1 or At1G53430. In addition, RIR1 and At1g53430 are expressed at similar protein levels while At1g53420 is extremely low abundant (Mergner et al., 2020); proteomicsDB: https://www.proteomicsdb.org). In analogy to RIR1, At1g53430 contains 11 LRRs followed by a malectin domain prior in the ectodomain (Fig. 7A). This RLK has been previously described to be transcriptionally induced during nematode infection and was therefore named NEMATODE-INDUCED LRR-RLK 2 (NILR2) (Mendy et al., 2017). However, a *nilr2* mutant did not exhibit any susceptibility phenotype upon infection with nematodes (Mendy et al., 2017).

**Fig. 6:**
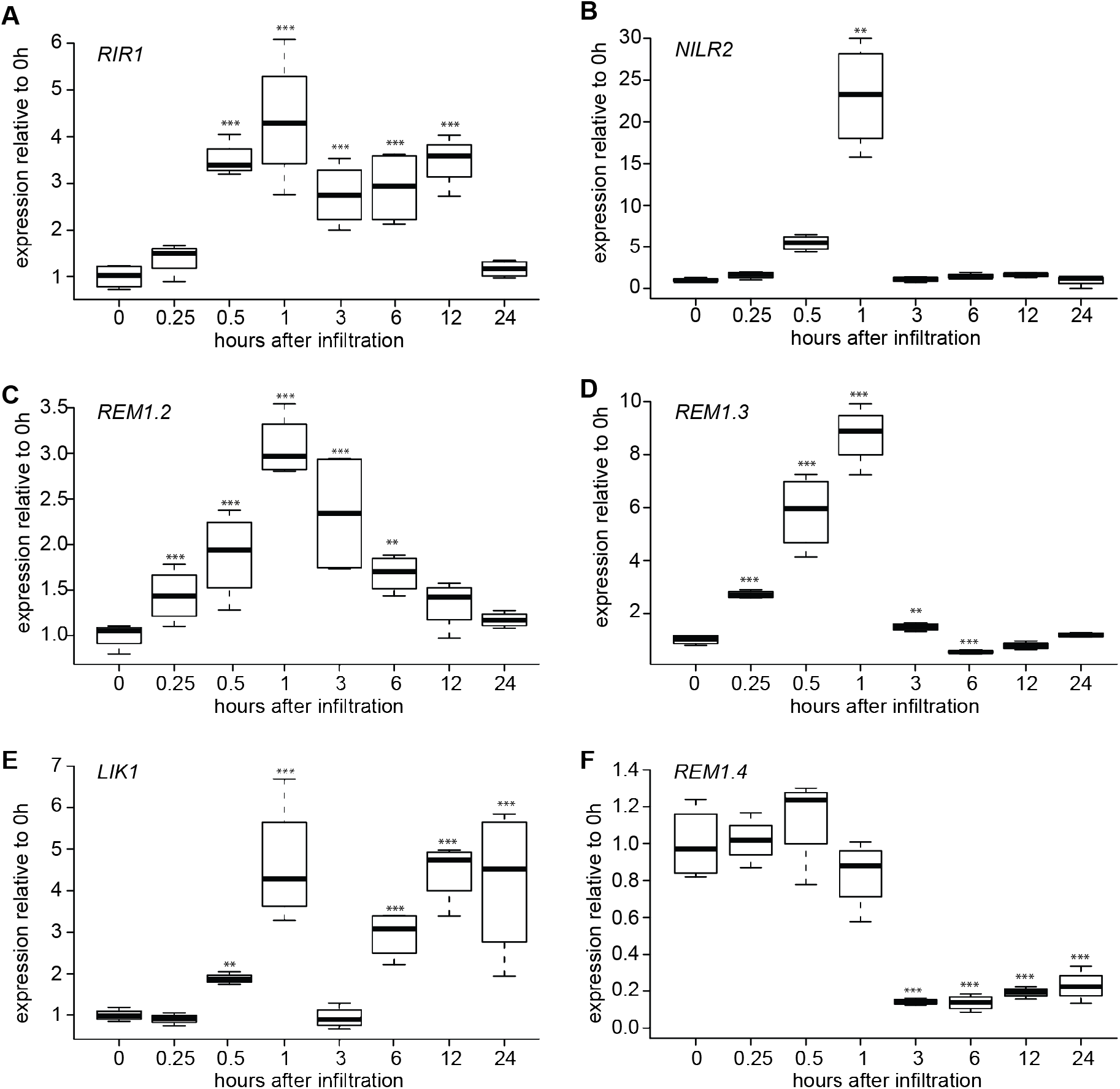
*RIR1, NILR2, REM1.2* and *REM1.3* are up-regulated upon water infiltration. Five-week-old Col-0 leaves were infiltrated with H_2_O and harvested at the indicated time points (n =4). *RIR1* (A), *NILR2* (B), *REM1.2* (C), *REM1.3* (D), *LIK1* (E) and *REM1.4* (F) transcripts were normalized to *UBC*. Transcripts in the non-infiltrated control (0 h) were set to 1, respectively. Statistical significance was assessed using one-way ANOVA followed by Dunnett’s multiple comparison test (** p≤ 0.01, *** p≤ 0.001).

**Fig. 7:**
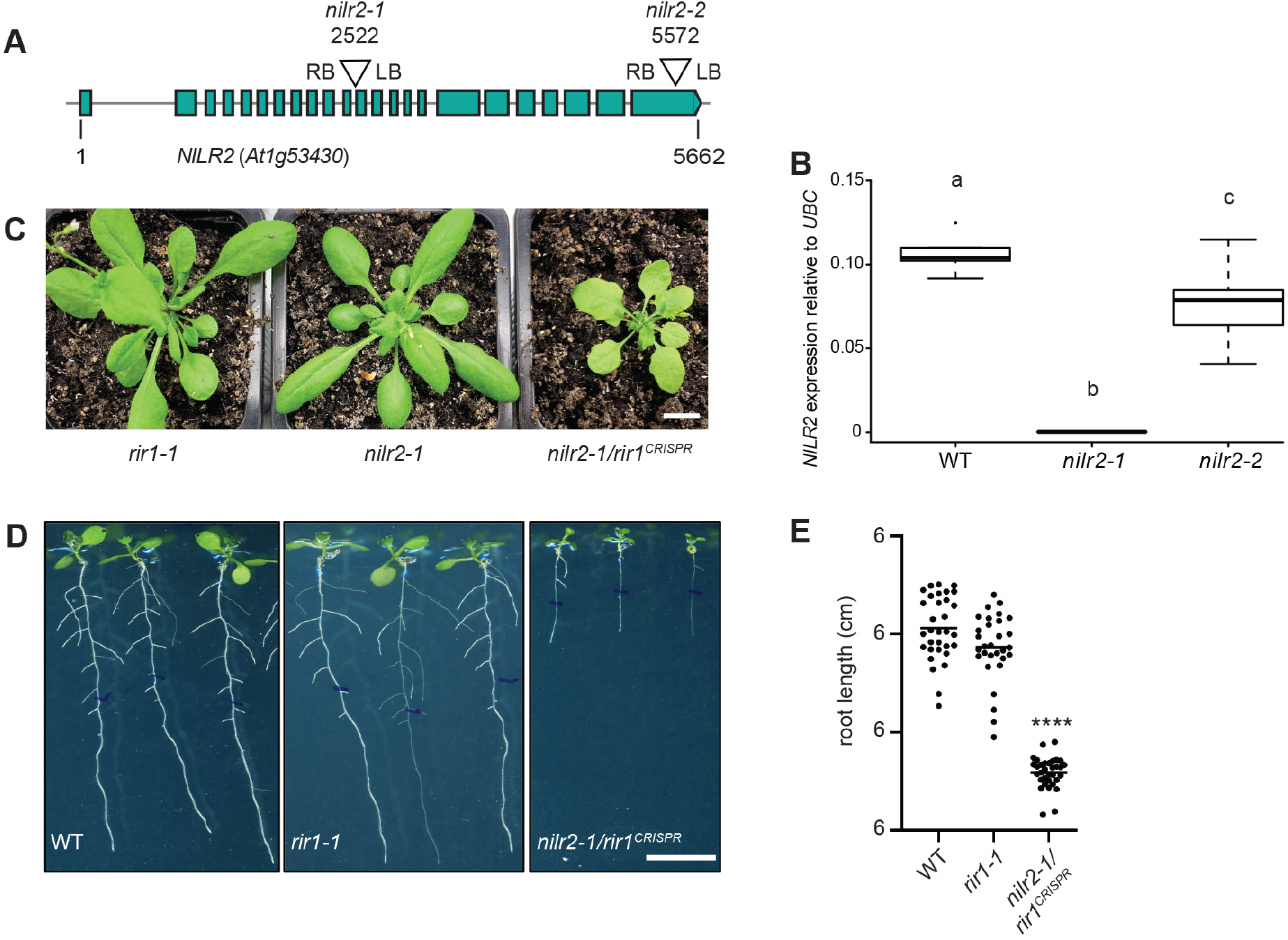
RIR1 and NILR2 form a functionally redundant gene cluster. (A) Schematic illustration of the *NILR2* gene (At1g53430) that is adjacent to the *RIR1* locus. (B) Quantitative RealTime PCR on *nilr2* alleles demonstrates *nilr2-1* being a genetic knock-out while *nilr2-2* transcripts were less strongly affected. Statistical significance for each transcript was assessed using ANOVA followed by Tukey HSD test (p<0.05). (C) While the *rir1-1* allele did not show any growth phenotype, a *rir1/nilr2* double mutant generated by introducing a CRISPR-CAS9-mediated 1725bp mutation in the *RIR1* locus (*nilr2-1/rir1^CRISPR^*) resulted in strongly retarded growth in seedlings. (D) In vitro growth phenotype scored on plants grown for ten days on MS agar plates. (E) Quantification of the *nilr2-1/rir1^CRISPR^* root phenotype. Statistical significance for each transcript was assessed using ANOVA followed by a Tukey multiple comparison test (p<0.0001). Scale bars indicate 1 cm.

To get more insight into a functional relationship between RIR1 and NILR2, we searched publicly available transcriptome datasets. Here, RIR1 and NILR2 transcripts were shown to be induced in Arabidopsis upon treatment of seedlings for 30 minutes with the bacterial elicitor flg22 (Bjornson et al., 2020), we tested transcript abundance of both receptors over a prolonged incubation period. For this, we syringe-infiltrated a 1 μM flg22 solution into leaves of 5-week-old Arabidopsis plants and harvested them after 1 and 24 hours. At neither time point transcript levels of *RIR1* and *NILR2* were different compared to the water control (Fig. S6). However, we observed a significant reduction between 1 and 24 hours of flg22 treatment as well as in the controls. Since this effect was also found in water-infiltrated plants, we scored transcriptional changes in a detailed time course experiment. Interestingly, infiltration of water was sufficient to transiently induce *RIR1* and *NILR2* expression, reaching a maximal expression at 1 hour post infiltration (Fig. 6A-B). As our data show that RIR1 directly interacts with the heterooligomeric REM1.2/1.3 complex, we also tested expression of these genes and found that *REM1.2* and *REM1.3* transcripts followed a similar pattern (Fig. 6C-D). However, expression patterns of the related malectin-domain receptor *LIK1*, that has previously been shown to interact with the cytokinin receptor CERK1 (Le et al., 2014) and the remorin *REM1.4* were distinct from *RIR1, NILR2, REM1.2* and *REM1.3* (Fig. 6E-F). To test whether these transcriptional inductions are an integral part of a wounding response triggered by water infiltration, we mechanically injured five weeks old rosette leaves with a needle and scored transcript abundance at one and 24 hours after wounding. No induction was found after one hour, receptor transcripts were found to be reduced (Fig. S7A-B) while REM1.2 and REM1.3 barely responded to the stimulus at all (Fig. S7C-D). In contrast, expression of the wounding marker genes *JASMONATE-ZIM-DOMAIN PROTEIN 10 (JAZ10;* At5g13220) and *LIPOXYGENASE 3 (LOX3;* At1g17420) were induced after one hour and reduced at 24 hours after wounding (Fig. S7E-F). These data indicate that the observed transcriptional inductions of *RIR1*, *NILR2*, *REM1.2* and *REM1.3* upon water infiltration do not represent a classical wounding response.

To test functional redundancy on a genetic level, we isolated two independent homozygous *nilr2* mutant alleles (*nilr2-1* (SALK_047602.23.65.n); *nilr2-2* (SALK_129312C)) (Fig. 7A). Transcript analysis demonstrated only *nilr2-1* to be a transcriptional null while *nilr2-2* transcripts were only moderately affected (Fig. 7B). However, and in analogy to our *rir1* mutants we did not observe any significant developmental phenotype in any of the two alleles (Fig. 7C).

Since chromosomal proximity of RIR1 and NILR2 did not allow the generation of double mutants by crossing, we introduced a CRISPR-CAS9 construct using four guide RNAs against RIR1 into the *nilr2-1* mutant background. Here guide RNAs 87 and 14 (see Table S4) were effective and generated a 1725bp deletion between bases 1938 and 3662 of the RIR1 genomic locus. Quantitative RealTime-RT-PCR showed the absence of transcript for both genes indicating these lines to be knock-down alleles (Fig. S8). Homozygous *nilr2-1/rir1^crispr^* double mutants were developmentally retarded (Fig. 7C-D) and thus phenocopied the *rem1.1/1.2/1.3/1.4* quadruple mutant during an early phase of development. However, this retardation was compensated over time leading to plants that were phenotypically indifferent from the WT upon flowering. These data suggest that RIR1 and NILR2 are functionally redundant. Since we were so far unable to unravel the precise molecular function of both receptors, further studies are needed to define the precise interplay and mode of action of these two receptors.

## Discussion

Although remorin proteins have been intensively studied in the recent past (reviewed in Gouguet et al., 2020) the precise molecular function remains elusive for most of them. However, some canonical features have been unraveled, among them their ability to from higher order oligomers (Bariola et al., 2004; Marin et al., 2012; Legrand et al., 2019; Martinez et al., 2019) and to interact with receptor-like kinases (Lefebvre et al., 2010; Gui et al., 2016; Liang et al., 2018). Among the remorin protein family, the phylogenetic Group 1 (Raffaele et al., 2007) stands out due to the high abundance of REM1.2 and REM1.3. However, it was only until recently that general growth defects have been observed for *rem1.2/1.3* double mutants (Huang et al., 2019). Although we could not confirm this phenotype under our growth conditions when using the same genetic material, a higher order quadruple mutant indeed exhibited a dwarfism phenotype (Fig. 1D). Since the severity of the phenotype may depend on the growth conditions and likely to environmental stresses, it is possible that such differences explain the discrepancy between phenotypical observations of the *rem1.2/1.3* double mutants. Our genetic and protein data further support the existence of an endogenous hetero-oligomeric remorin complex in *A. thaliana* (Fig. 2A). Both, REM1.2 and REM1.3 have a largely overlapping interactome, although they can still be clearly distinguished from each other using PCA analysis (Fig. 2B-C). Taken together these data indicate that Group 1 remorins form a hetero-oligomeric scaffold at the plasma membrane that serves core functions in plant growth and development and in responses to stress (e.g. salt stress and viral infections; Fig. 1).

The involvement of remorins in general plant growth by restricting plasmodesmal aperture as previously suggested (Perraki et al., 2014; Gronnier et al., 2017; Huang et al., 2019) was confirmed using Arabidopsis double mutants (Fig. 1B). The growth retarded phenotype of the *rir1/nilr2* double mutant during early seedling development (Fig. 7) places both receptors into this genetic pathway, although a precise role remains to be unraveled. The genetic interaction between RIR1 and NILR2 is also supported by a recent in silico receptor interaction network analysis that indicated an association of RIR1, NILR2 and a so far uncharacterized LRR-RLK (At5g01950) (Xi et al., 2019). Interestingly, the phylogentically related MD-receptor-like kinase LMK1 (At1g07650) was reported to function as a cell death-related receptor as it induced cell death when ectopically expressed in *N. benthamiana* leaf epidermal cells (Li et al., 2020). While we did not observe such phenomenon for RIR1, we were indeed also unable to over-express NILR2 in these cell types (data not shown). Possible detrimental effects due to ectopic expression in their homologous plant backgrounds have also been reported for Arabidopsis (Huang et al., 2019); Fig. S1B) and *Solanum lycopersicum* (Cai et al., 2020) remorin proteins. In addition, we have not succeeded generating stable transgenic Arabidopsis lines constitutively over-expressing group 1 remorins at high level.

The presence of an MD in both receptors indicates possible functions in sensing cell wall integrity as an animal malectin protein has been shown to bind Glc2-N-glycans (Schallus et al., 2008; Schallus et al., 2010). Besides MDs, some plant RLKs like the CrRLK1Ls contain two malectin-like domains (MLD) in a tandem organization in their extracellular region (Boisson-Dernier et al., 2011). Members of this family are thought to bind to the cell wall and act as sensors of cell wall integrity (Cheung and Wu, 2011). In line with this, several CrRLK1Ls have been shown to be involved in various growth-and stress-related pathways (Galindo-Trigo et al., 2016). Mutations in the RLK THESEUS1 (THE1), for example, were demonstrated to partially restore growth defects of the cellulose-deficient mutant *prc1-1*, without altering the cellulose content (Hématy et al., 2007). In addition, a functional THE1 is needed for the oxidative burst triggered by the cellulose synthesis inhibitor isoxaben (Denness et al., 2011). However, it was recently demonstrated that MLDs can bind RALF peptides and that they lack key residues required for direct carbohydrate binding (Xiao et al., 2019).

Since a huge variety of environmental stimuli needs to be integrated simultaneously and given the limited capability of lateral diffusion of membrane proteins in plant cells (Martinière et al., 2019), scaffolding proteins could dynamically facilitate and compartmentalize these processes. However, RIR1 mobility and its recruitment to membrane nanodomains is not altered in remorin mutants (Fig. S4), indicating REM1.2 and REM1.3 do not regulate lateral diffusion of RIR1 at the PM itself. This is in contrast to the group 2 remorin SYMREM1 from *Medicago truncatula* that immobilizes the entry receptor LYK3 in membrane nanodomains during symbiotic interaction with rhizobia (Liang et al., 2018). However, it was recently shown that the presence of Group 1 remorins can alter membrane fluidity (Huang et al., 2019) and as a consequence the dynamic behavior of adjacent membrane proteins. Whether this is also the case for FLS2 that has been shown to partially co-localize in REM1.2/REM1.3 positive membrane nanodomains (Bücherl et al., 2017) remains to be demonstrated. However, we hypothesize at this stage that the transient transcriptional induction of Group 1 remorins upon infiltration of water into leaves (Fig. 7) may be part of a cellular response to minimize membrane damage under these conditions. It would indeed be interesting to test, whether the ability of remorins to form highly oligomeric proteinaceous leaflets (Martinez et al., 2018) or potentially liquid-liquid-phase separates (LLPS) (Jaillais and Ott, 2019) may assist membrane stabilization under conditions of negative curvature and membrane tension that cannot be compensated by the rigid cell wall. Given the presence of the malectin domain in RIR1 and NILR2 (Fig. 3B, 7A), these receptors might act as sensors of PM-cell wall proximity and get activated upon retraction of the PM from the cell wall.

## Material and Methods

### Plant materials and growth conditions

T-DNA insertion mutants of REM1.2 (At3g61260) *rem1.2-2* (SALK_117637.50.50.x), REM1.3 (At2g45820) *rem1.3-2* (SALK_117448.53.95.x) and REM1.4 (At5g23750.x) *rem1.4-3* (SALK_073841.47.35), *rir1-1* (Salk_130548.42.45.x), *rir1-2* (Salk_057812.14.90.x) were provided by the ABRC. *rem1.2-2/rem1.3-2* (*rem1.2/rem1.3*) double mutant was generated by crossing the respective T-DNA inserted parental plants, *rem1.2-2/rem1.3-2/rem1.4-3* (*rem1.2/rem1.3/rem1.4*) was created by crossing *rem1.2/rem1.3* with *rem1.4-3*. All plants were genotyped using primers indicated in Table S4.

ProREM1.2:YFP-REM1.2 and ProREM1.3:YFP-REM1.3 lines were described previously (Jarsch et al., 2014).

Surface-sterilized *A. thaliana* seeds were germinated on ½ MS plates supplemented with 1% sucrose. If not indicated differently seedlings were transferred to soil and grown under short day conditions (8 h light/16 h dark). For immunoprecipitation followed by mass spectrometry, seeds of indicated lines were sterilized and germinated on metal grids placed on solid ½ MS media and grown for 1 week. Metal grid including germinated seedlings were moved to liquid ½ MS media and grown for 3 weeks under long day conditions, liquid media was replaced with fresh media every week.

The *ProRIR1:RIR1-GFP* construct (in pGWB604) was transformed into *rir1-1, rem1.2-1* and *rem1.2-2/rem1.3-2* mutant plants by the floral dipping method ((Clough and Bent 1998)). The *ProREM1.2-mCherry-gREM1.2* and *ProREM1.2-mCherry-gREM1.3* constructs (in pGWB1) were additionally dipped into homozygous *ProRIR1:RIR1-GFP/rir1-1* plants.

*Nicotiana benthamiana* plants were grown for 4 weeks under long day conditions (16 h light/8 h dark) prior to *Agrobacterium*-mediated transformation.

### Bacteria strains

All cloning was performed using *E. coli* Top10 strains employing standard protocols for transformation, cultivation and lysis. Electro competent *Agrobacterium tumefaciens* AGL1 was used for Arabidopsis transformation.

### Developmental phenotyping

Seedlings designated for examination of germinations rates and rosette diameter at 14 days were grown on vertical plates under long day conditions. The phenotyping of later development stages was performed on adult plant grown under long day conditions in the greenhouse.

### Salt stress phenotyping

Seeds of indicated genotypes were sterilized then germinated on ½ MS plates and grown for 1 week before transfer to new plates including 100 mM NaCl or mock, seedlings were grown for 1 week on NaCl plates before root elongation was measured.

### Viral spreading experiments

PlAMV propagation experiments were performed as previously described (Gronnier et al., 2017) with minor modifications. *Agrobacterium tumefaciens* strain GV3101 carrying pLI1evec-nGFP, which expresses PlAMV-GFP (Minato et al., 2014) was infiltrated on 4-week-old *Arabidopisis thaliana* plants grown in a greenhouse, with a final OD600 nm=0.2. Viral spreading was tracked on inoculated leaves using Axiozoom macroscope system (Zeiss, Germany) at 7 DPI. The area of PlAMV-GFP infection foci was measured using Fiji software (http://www.fiji.sc/).

### Quantitative real-time PCR

RNA from 10 days old seedlings were extracted using a Spectrum Plant Total RNA Kit (Sigma). DNA was removed by DNase I Amplification Grade (Invitrogen). cDNA was generated using a qScript cDNA SuperMix (Quanta bio). qPCR was performed with a Fast SYBR kit (Applied Biosystems) on a StepOnePlus Real-Time PCR system machine (Applied Biosystems) or a 96-well Real-Time PCR machine (Bio-Rad, CFX96).

Two technical and at least 3 biological replicates were used. All transcripts were normalized to the housekeeping gene *AtUbiquitin C (UBC*). All primers used can be found in Supplemental Table S4.

For assessing gene expression under different conditions, five-week-old Arabidopsis leaves were used. Leaves were infiltrated with water or 1 μM flg22 and water (as control) for the examination of the defense marker genes or wounded with a needle to analyze the stress induction. Here, non-infiltrated and non-wounded leaves were used as controls.

### Yeast two-hybrid assay

Yeast two-hybrid interaction assays and expression analyses were performed as previously described (Marin et al., 2012). In brief, using the vectors pGADT7:GW and pGBKT7:GW, activation domain (AD)- and binding domain (BD)-fused constructs were co-transformed into the yeast strain PJ69-4a, respectively. Transformants were plated onto different SD plates lacking tryptophan, leucine and histidine (+/-3-amino-1,2,4-triazole) to examine the interaction. The expression of all proteins was verified by Western Blot analysis using tag-specific antibodies.

### Confocal laser-scanning microscopy

For microscopic analyses, *A. thaliana* leaves were infiltrated with H_2_O prior to harvest (puncher 4 mm diameter). Leaf discs were mounted on glass slides and immediately imaged using Leica TCS SP5 and SP8 confocal microscopes. GFP was excited with the Argon laser (AR) at 488 nm and the emission was detected between 500 and 550 nm. YFP RICKY1 was excited with a wavelength of 514 nm (AR) and the emission was detected at 525 – 599 nm. mCherry was excited at 561 nm using the Diode Pumped Solid State (DPSS) laser and the emission was detected at 570 – 640 nm. Samples, co-expressing two fluorophores, were imaged using the sequential mode between frames.

### Fluorescence lifetime imaging (FLIM)

Cotyledons of 5 days old Arabidopsis seedlings (*rir1-1/ProRIR1:RIR1-GFP, rir1-1/ProRIR1:RIR1-GFP/ProREM1.2:mCherry-gREM1.2* and *rir1-1/ProRIR1: RIR1-GFP/ProREM1.3:mCherry-gREM1.3*) were used for FLIM experiments. Experiments were performed using a Leica TCS SP8X confocal microscope equipped with TCSPC (time correlated single photon counting) electronics (PicoHarp 300), photon sensitive detectors (HyD SMD detector) and a pulsed white light laser. The abaxial side of cotyledons were imaged using a 63x/1.2 water immersion objective lens (Leica C-APOCHROMAT 63×/1.2 water). GFP was excited at 488 nm and fluorescence emission collected between 500-540nm. The FLIM data sets were recorded using the Leica LASX FLIM wizard linked to the PicoQuant SymPhoTime 64 software and acquired by repetitively scanning (40 times) region of 70.29 μm2 at 50Mhz with a pixel dwell time of 9.75 μs. After each FLIM acquisition effective expression of REM1.2 and REM1.3 was recorded by exciting mCherry at 580 nm and collection of corresponding fluorescence emission between 590 and 640 nm. To visualize potential FRET, samples were excited at 488 nm and fluorescence emission was collected between 590 and 640 nm. The instrument response function (IRF) was measured using erythrosine B as described (Stahl et al., 2013). Calculations of fluorescence lifetime were performed using the PicoQuant SymPhoTime 64 software following instructions for FLIM-FRET-Calculation for Multi-Exponential Donors. A two-exponential decay fit was used for GFP. The lifetimes were initially estimated by fitting the data using the Monte Carlo method and then by fitting the data using the Maximum Likely Hood Estimation.

### Total Internal Reflection Fluorescence (TIRF) microscopy

TIRF microscopy was performed using a Zeiss Elyra PS1 (Zeiss, Germany) microscope using a 100 × 1.4 NA oil immersion objective. GFP was excited using a 488 nm solid-state laser diode and corresponding fluorescence emission was collected with an EM-CCD camera with bandwidth filters ranging from 495–550 nm. The optimum critical angle was determined as giving the best signal-to-noise. Images time series were recorded at 10 frames per second (100 ms exposure time). To analyze single particle tracking experiments, we used the plugin TrackMate 2.7.4 (Tinevez et al., 2016) in Fiji (Schindelin et al., 2012) as previously described (Gronnier et al., 2020).

### Total protein extraction

Tissue from Arabidopsis were harvested in equal amount and frozen in liquid nitrogen before grounded to power by a tissue lyser (for small amount) or mortar and pestle (for larger amounts). Proteins were extracted in a buffer containing: 50 mM Tris-HCl (pH 7.5), 150 mM NaCl, 10 % glycerol, 2 mM EDTA 5 mM DTT, 1 mM PMSF in 2-propanol, 1.5 mM Na3VO4, 1 x Protease Inhibitor Cocktail (Roche) and 1 % (v/v) IGEPAL CA-630 (Sigma-Aldrich). Extraction was incubated for 1 hour at 4°C on a rotor. After incubation extracts were centrifuged at 13,000 rpm for 30 min. Supernatants were mixed with SDS loading buffer and boiled for 5 min.

### Immuno-precipitation mass spectrometry

For determining remorin interactions, tissue from Arabidopsis were harvested in equal amount and frozen in liquid nitrogen before grounded to power by a tissue lyser. Proteins were extracted in a buffer containing: 50 mM Tris-HCl (pH 7.5), 150 mM NaCl, 10 % glycerol, 2 mM EDTA 5 mM DTT, 1 mM PMSF in 2-propanol, 1.5 mM Na3VO4, 1 x Protease Inhibitor Cocktail (Roche) and 1 % (v/v) IGEPAL CA-630 (Sigma-Aldrich). Extraction was performed for 1 hour at 4°C on a rotor. After incubation extracts were centrifuged at 13,000 rpm for 30 min. Supernatants were incubated with GFP-Trap (Chromotek) for 2 hours at 4°C on a rotor before magnetic separation and washing 5 times with washing buffer (50 mM Tris-HCl (pH 7.5), 150 mM NaCl, 10 % glycerol, 2 mM EDTA 5 mM, 1 x Protease Inhibitor Cocktail (Roche) and 0.5 % (v/v) IGEPAL CA-630 (Sigma-Aldrich)). Extracted proteins were released with 2xNuPage LDS sample buffer and boiled for 5 min.

Samples were reduced with 10 mM ditiothreitol (DTT) for one hour followed by alkylation with 55 mM chloroacetamide (CAA) for 30min at room temperature and run into a 4-12% NuPAGE gel (Invitrogen) for approximately 1 cm. In-gel digestion was performed according to standard procedures with trypsin (Roche) (Shevchenko et al., 2006). MS measurements were performed on a Q Exactive HF-X (Thermo Fisher Scientific) using a 60 min linear gradient. The instrument was operated in data-dependent mode, automatically switching between MS and MS2 scans. Full-scan mass spectra (m/z 360-1300) were acquired in profile mode with 60,000 resolution, an automatic gain control (AGC) target value of 3e6 and 45 ms maximum injection time. For the top 20 precursor ions, high resolution MS2 scans were performed using HCD fragmentation with 28 % normalized collision energy, 30,000 resolution, an AGC target value of 2e5, 50 ms maximum injection time and 1.3 *m/z* isolation width in centroid mode. The minimum AGC target value was set to 2.2e3 with a dynamic exclusion of 25s.

Peptide and protein identification and quantification was performed with MaxQuant (Cox and Mann, 2008) using standard settings (version 1.6.3.3). Raw files were searched against an Arabidopsis database (Araport11_genes.201606.pep.fasta) and common contaminants. Carbamidomethylated cysteine was set as fixed modification and oxidation of methionine, and N-terminal protein acetylation as variable modifications. Trypsin/P was specified as the proteolytic enzyme, with up to two missed cleavage sites allowed. The match between run and LFQ functions were enabled and results filtered to 1% PSM and protein FDR. LFQ proteinGroup intensities were filtered for identifications in all three replicates of at least one group (n = 2,195), missing values were imputed from a normal distribution and statistical analyses were performed using Perseus (v. 1.6.0.2) (Tyanova et al., 2016).

To assess RIR1 interactions, extracts of 5- to 6-week-old *A. thaliana* plants were incubated with 50-100 μl previously washed 50% GFP-binding protein beads slurry (Chromotek, GFP-Trap^®^) at 4 °C for 3 hours on a roller mixer. Samples were centrifuged to collect beads; supernatants were discarded and the beads were washed 5 times using extraction buffer containing 0.5 % (v/v) IGEPAL CA-630 (Sigma-Aldrich). To release the proteins, 50-100 μl 2x NuPAGE LDS sample buffer (Invitrogen) was added and samples were boiled for 5 min at 95 °C.

Obtained proteins were separated by SDS-PAGE (NuPAGE, Invitrogen) and after staining with Coomassie brilliant Blue G- ds were excised from the gel and transferred to LoBind tubes (Eppendorf). For in-gel digestion, gel pieces were washed twice for 30 min with Acetonitrile/Ammonium Bicarbonate (ACN/ABC) at 56 °C and dehydrated for 5 min using 100 % Acetonitrile at room temperature. Supernatants were discarded and di-sulphide bonds were reduced using 10 mM DTT in ABC for 45 min at 56 °C. For alkylation, samples were incubated for 30 min with 55 mM Chloroacetamide in ABC in the dark. Samples were again washed and dehydrated as described before. In-gel digest was performed with trypsin (Promega) in 50 mM ABC and 5 % ACN at 37 °C overnight. Peptides were recovered with 5 % Formic acid in 50 % ACN by repeated sonication. Supernatants were collected in new LoBind tubes and dried completely in speed vacuum for 3 hours at room temperature. LC-MS/MS analysis was performed using an Orbitrap Fusion mass-spectrometer (Thermo Scientific). The entire TAIR10 database was searched (www.Arabidopsis.org) using Mascot (v 2.3.02, Matrix Science) with the inclusion of sequences of common contaminants, such as keratins and trypsin. Parameters were set for 5 ppm peptide mass tolerance and allowing for Met oxidation and two missed tryptic cleavages. Carbamidomethylation of Cys residues was specified as a fixed modification, and oxidized Met and phosphorylation of Ser, Tyr or Thr residues were allowed as variable modifications. Scaffold (v3; Proteome Software) was used to validate MS/MS-based peptide and protein identifications and annotate spectra.

### SDS-PAGE and Western blots

Extracted proteins were separated on a 12% SDS-PAGE gel for 15 min at 80V followed by 2-3H at 120V. Separated proteins were transferred to PVDF membranes at 30V for 12-16H at 4°C. PVDF membranes were blocked with 5% milk followed by an incubation with first antibodies for REM1.2/1.3 followed by 3 washes and an incubation with second α-rabbit antibody fused to HRP against rabbit (Sigma). Clarity Western 701 ECL (Bio-Rad) was employed for Chemiluminescence detection.

## Acknowledgments

This project was supported by the German Research Foundation (Deutsche Forschungsgemeinschaft; DFG) in frame of the Collaborative Research Center 924 (SFB 924; T.O.), under Germany’s Excellence Strategy/Initiative (CIBSS – EXC-2189 – Project ID 390939984; T.O.), supplemented by an Exploration Grant of the Boehringer Ingelheim Foundation (to T.O.), an individual DFG grant KE1485 (to B.K.), the Gatsby Charitable Foundation (C.Z.), the University of Zürich (C.Z.), the European Research Council under the Grant Agreements 309858 and 773153 (grants PHOSPHinnATE and IMMUNO-PEPTALK to C.Z.) and the Swiss National Science Foundation (grant no. 31003A_182625 to C.Z.). J.G. was supported by the European Molecular Biology Organization (EMBO Long-Term Fellowship 438-2018). We thank the staff of the Life Imaging Center (LIC) in the Center for Biological Systems Analysis (ZBSA) of the Albert-Ludwigs-University Freiburg for help with their confocal microscopy resources, and the excellent support in image recording. The CLSM Leica SP8 was funded by the DFG grant INST 39/1104-1 FUGG.

Furthermore, we would like to thank Johannes Stuttmann for discussions and for providing the CRISPR-Cas9 vectors, Eija Schulze for the great technical assistance, Elisabeth Lichtenberg and Jacqueline Cornean for their help and all members of our team for fruitful discussions and providing their individual expertise throughout the course of the project. J.G. is also thankful to Grant Calder (JIC Bioimaging facility) for assistance in setting up FLIM-FRET experiments. GFP-PlAMV viral spreading was performed at the Bordeaux Imaging Center, member of infrastructure France Bioimaging.

## Supplemental Figures

**Figure S1:**
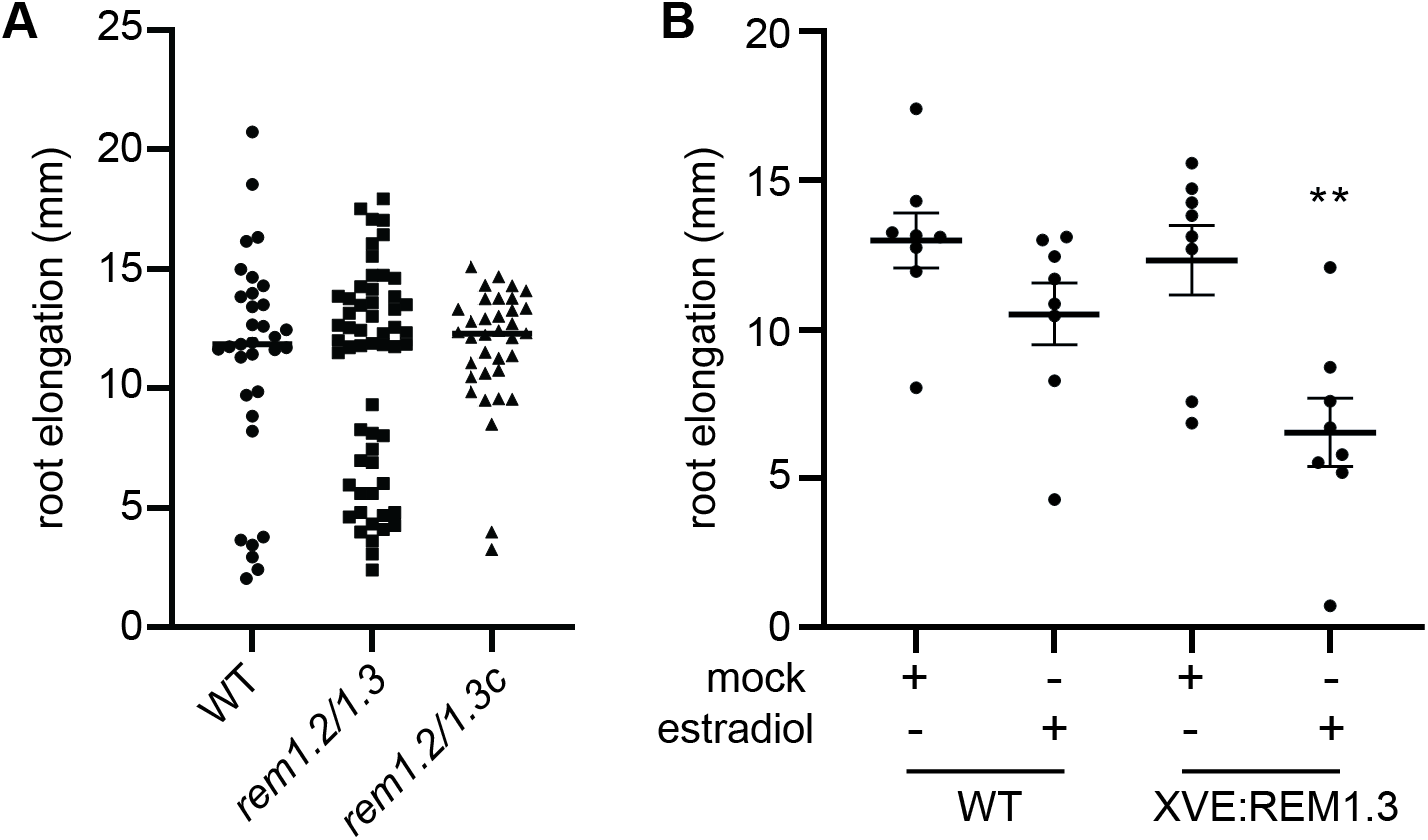
Assessing the growth phenotype of published remorin mutants under our growth conditions. (A) Root elongation of WT, *rem1.2/rem1.3* and *rem1.2/rem1.3c* (mutant published in Huang *et al*., 2019) over seven days. (B) Root elongation of WT and the published estradiol-inducible REM1.3 line XVE:REM1.3 three days after estradiol or mock treatment. Significance was tested using a Dunnett’s multiple comparisons test with p<0.01 (**).

**Fig. S2:**
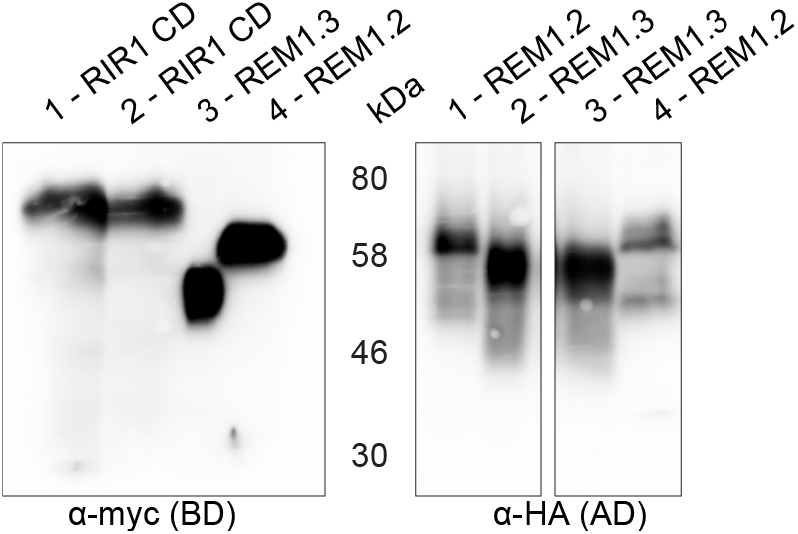
(B) Expression analysis of yeast clones using immunoblot analysis. Activation domain (AD)-fused proteins were visualized with an αHA antibody and binding domain (BD)-fused proteins with a α-myc antibody. CD= cytoplasmatic domain.

**Fig. S3:**
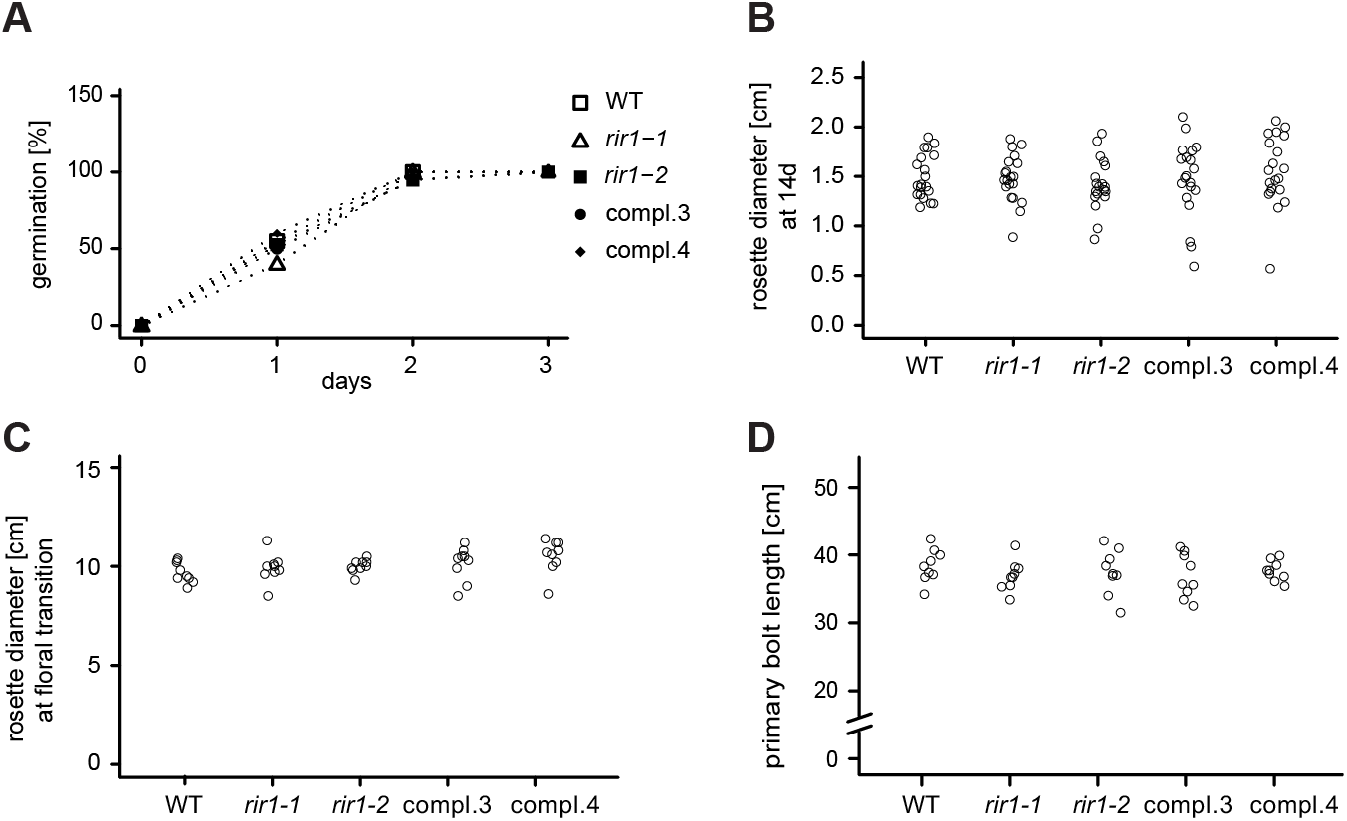
*rir1* mutants do not exhibit developmental defects. Different developmental stages were examined in Col-0 and *rir1* mutants grown under long-day conditions on plate (½ MS + 1 % sucrose) (A,B; n = 20) and on soil (C-F; n = 9). (A) Germination rates were scored on the indicated days. (B,C). The diameter of rosettes was measured in 14-day-old seedlings (B) and adult plants at the time of floral transition (C). The primary bolt length (D) was analyzed in seven-week-old plants. There was no significant difference between WT and the *rir1* mutants. Statistical significance was assessed using Fisher’s exact test (A) or one-way ANOVA (B-D), respectively.

**Fig. S4:**
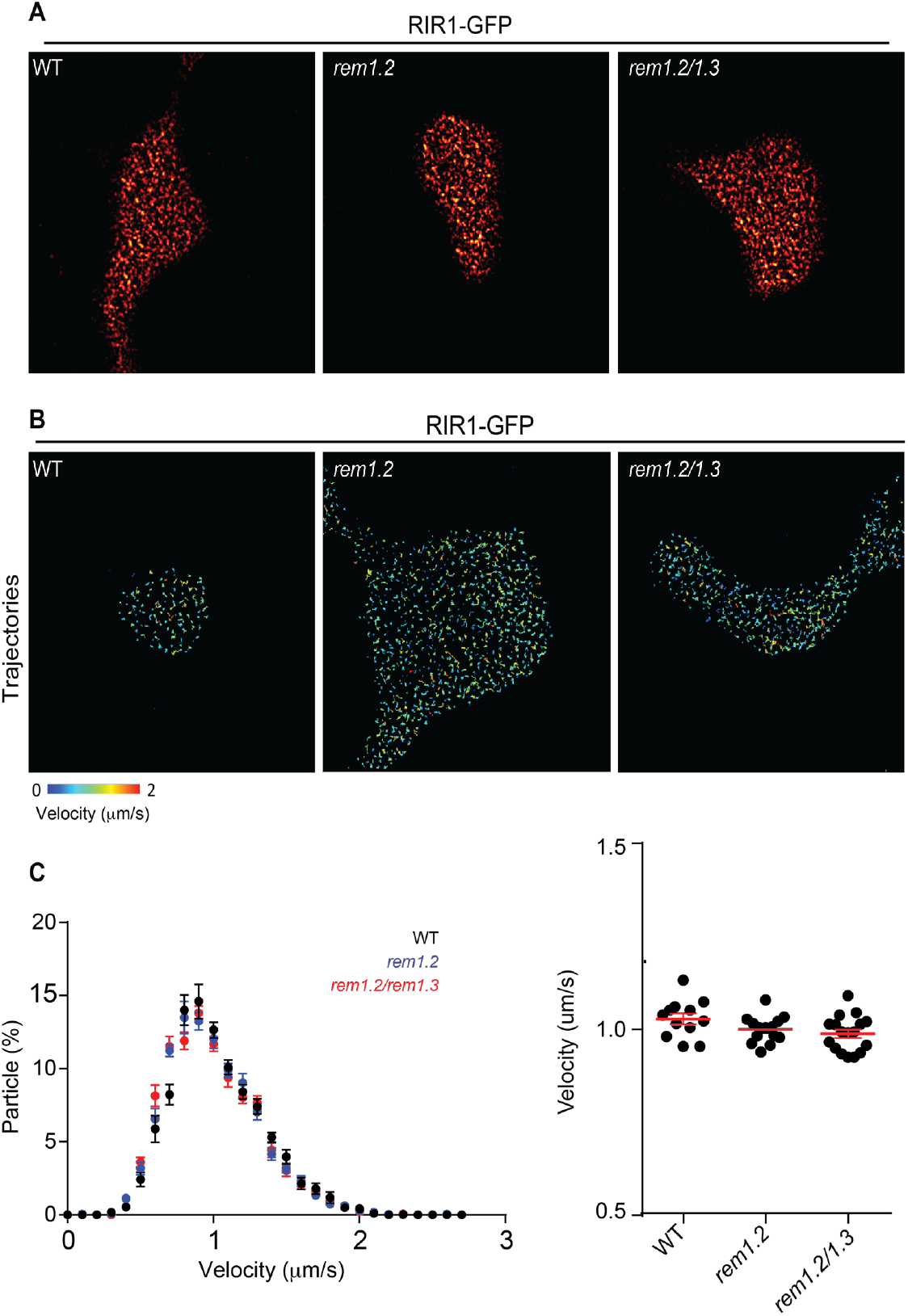
RIR1-positive nanodomains form in a REM1.2/1.3-independent manner. (A) TIRFM on *rir1*/RIR1-GFP, *rem1.2*/RIR1-GFP and *rem1.2/rem1.3*/RIR1-GFP. Experiments were performed on cotyledons of 5 days old seedlings. Image rendering: subtract background 20 rolling pixel and smoothing. (B) Single-particle tracking on *rir1*/RIR1-GFP, *rem1.2*/RIR1-GFP and *rem1.2/rem1.3*/RIR1-GFP. Experiment performed on cotyledons of 5 days old seedlings. (C) Quantification of RIR1-GFP single particle velocity. Mobility of RIR1-GFP is not altered in *rem1.2* or *rem1.2/rem1.3* mutants.

**Fig. S5:**
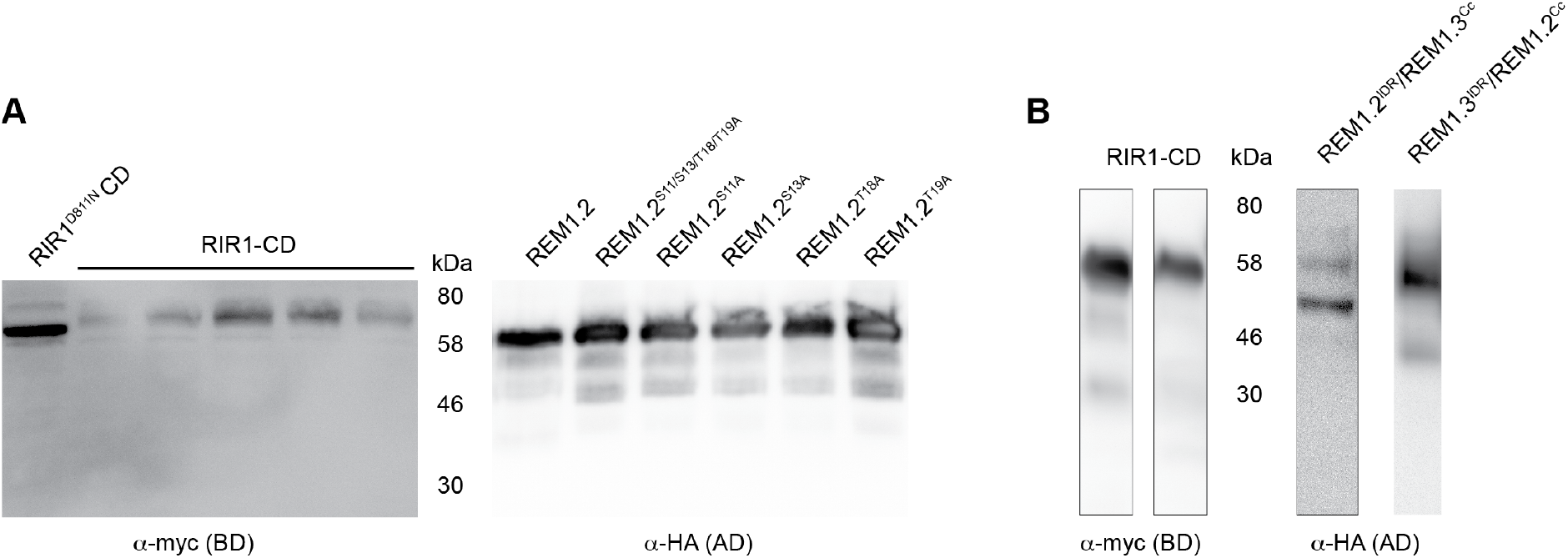
Western Blot analysis confirming the expression of clones that did not result in any interaction. (A) Blot supporting protein accumulations for Fig. 5A. (B) Blot supporting protein accumulations for Fig. 5B. Activation domain (AD)-fused proteins were visualized with an α-HA antibody and binding domain (BD)-fused proteins with an α-myc antibody. CD= cytoplasmic domain. Molecular weight is indicated in kDa.

**Fig. S6:**
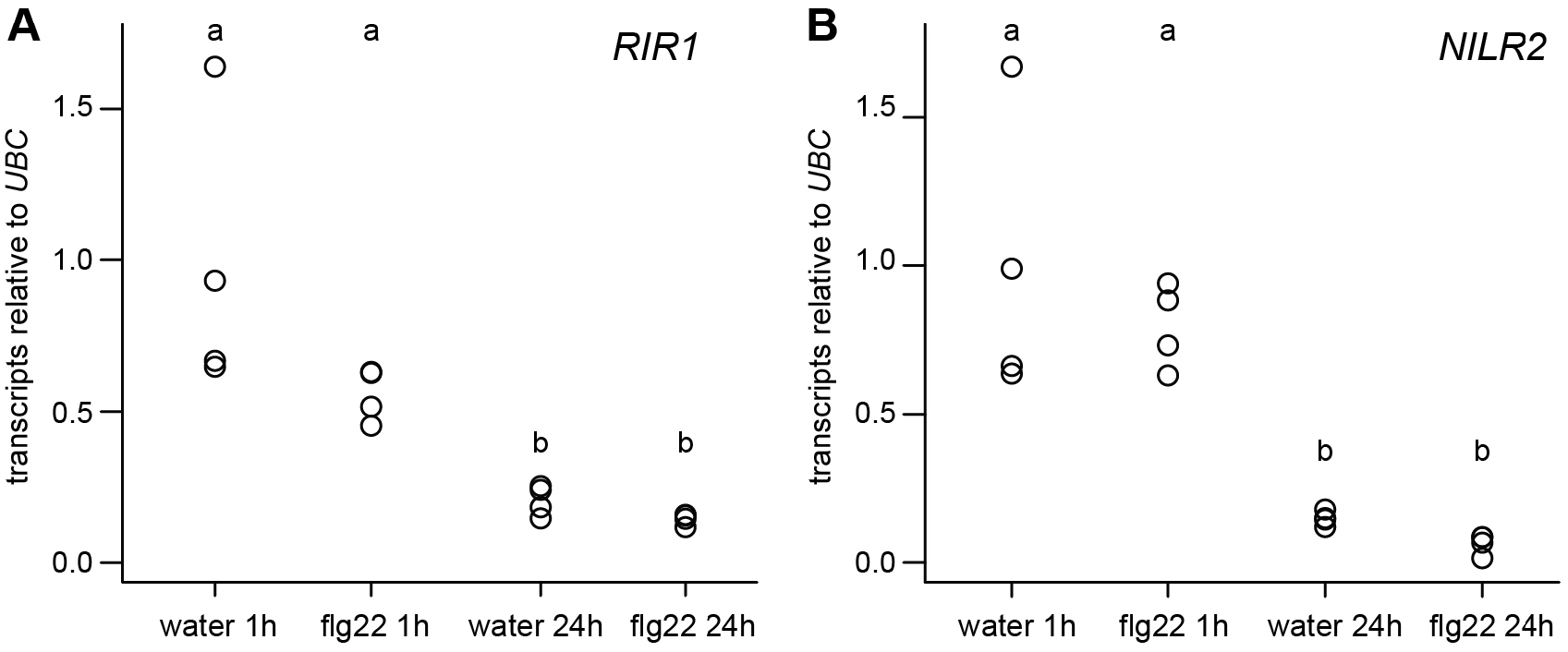
*RIR1* and *NILR2* transcripts are differently regulated 1 h and 24 h after infiltration. Five-week-old Col-0 plants were infiltrated with H_2_O or flg22 and harvested 1 h and 24 h later. *RIR1* (A) and *NILR2* (B) transcripts were analyzed using qPCR. Both transcripts were significantly lower 24 h after infiltration compared to 1 h after infiltration independent of the treatment. Statistical significance was assessed using one-way ANOVA followed by a Tukey HSD test (p≤ 0.005).

**Fig. S7:**
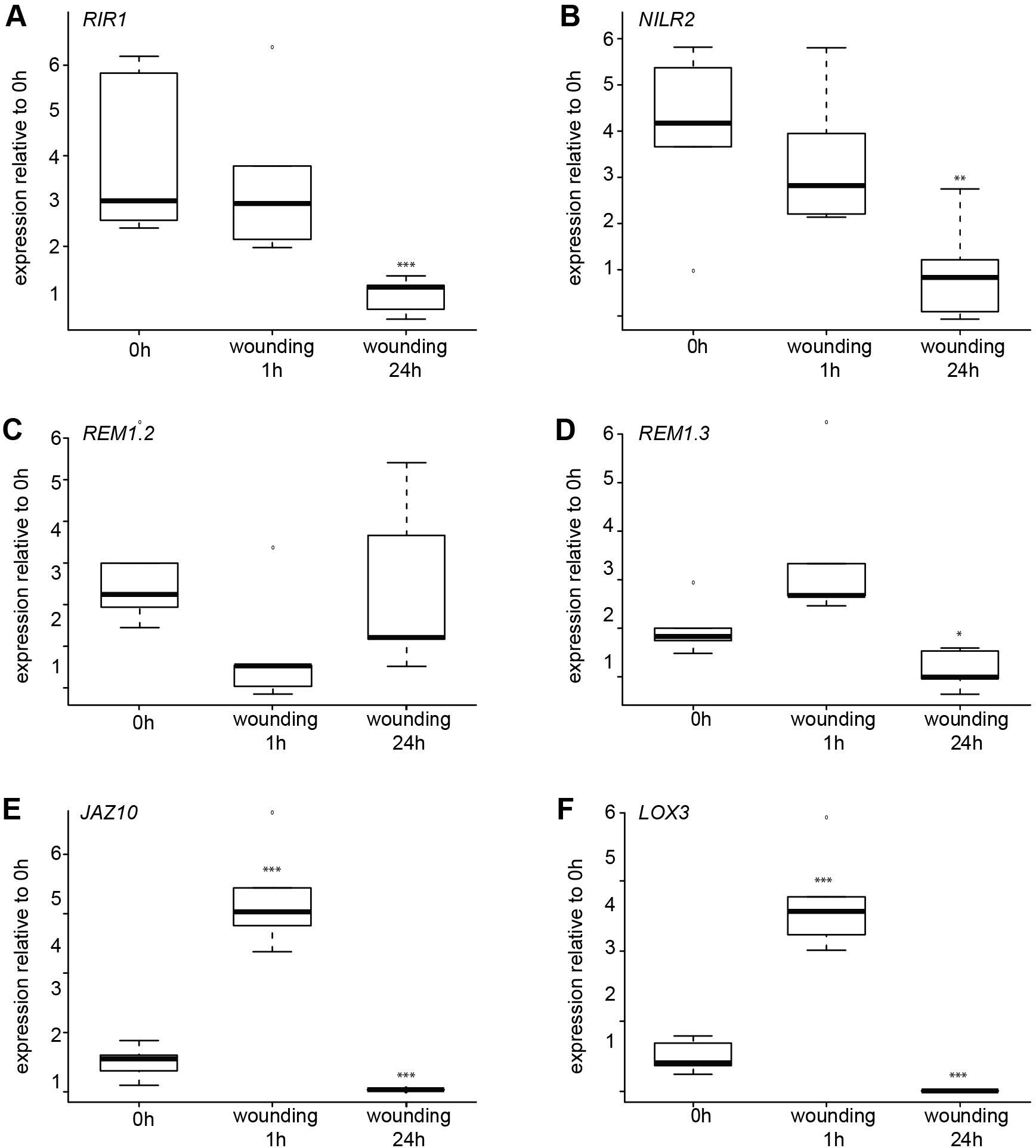
*RIR1, NILR2, REM1.2* and *REM1.3* transcripts are not up-regulated upon wounding. Five-week-old WT Col-0 leaves were wounded with a needle and harvested at the indicated time points. *RIR1* (A), *NILR2* (B), *REM1.2* (C), *REM1.3* (D), *JAZ10* (E) and *LOX3* (F) transcripts were normalized to UBC. The wounding marker genes *JAZ10* and *LOX3* were used as controls. Transcripts in the non-wounded control (0 h) were set to 1, respectively. Statistical significance was assessed using one-way ANOVA followed by Dunnett’s multiple comparison test (* p≤ 0.05, ** p≤ 0.01, *** p≤0.001).

**Fig. S8:**
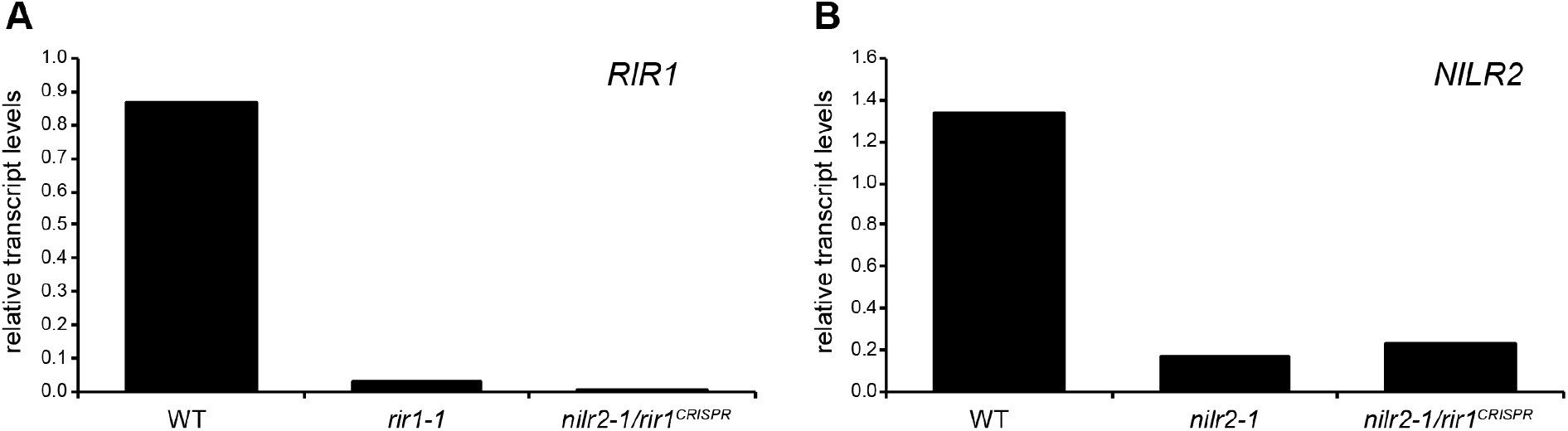
CRISPR-CAS9-mediated mutation of the *RIR1* locus in the *nilr2-1* mutant background. (A) Transcripts of *RIR1* in the *rir1-1* and the *nilr2-1/rir1^CRISPR^* mutant background. (B) Transcripts of *NILR2* in the *nilr2-1* and the *nilr2-1/rir1^CRISPR^* mutant background. Expression was determined by quantitative RealTime PCR and normalized to Ubiquitin.

